# Systematic CRISPRi screening reveals genetic modulators of *E. coli* isoprenoid production

**DOI:** 10.64898/2026.04.10.717724

**Authors:** Dhiraj Dokwal, Philip M. Brown, Christine Ingle, Scott H. Saunders, Kimberly A. Reynolds

**Affiliations:** Green Center for Systems Biology - Lyda Hill Department of Bioinformatics, The University of Texas Southwestern Medical Center, Dallas, TX 75230; Department of Biophysics, The University of Texas Southwestern Medical Center, Dallas, TX 75230

**Keywords:** Isoprenoid biosynthesis, CRISPR interference (CRISPRi), functional genomics, metabolic rebalancing, systems metabolic engineering, lycopene

## Abstract

Isoprenoid biosynthesis in *E. coli* is a promising source of high-value natural products. However, isoprenoid production imposes a substantial metabolic burden on cellular carbon and cofactor metabolism. Optimizing yields thus requires rebalancing gene expression throughout metabolism. To systematically identify the genes in *E. coli* central metabolism shaping isoprenoid yield, we established a high-throughput CRISPR interference (CRISPRi) assay to examine the impact of gene repression on production of the bright red carotenoid, lycopene. Our approach enables repression of both essential and non-essential genes and links gene expression perturbations to pathway yield and biomass. We screened a CRISPRi library targeting 180 *E. coli* genes and found 31 genes for which repression significantly modified lycopene yield. Genes whose repression increased lycopene yield were found across fatty acid, amino acid, central carbon, and isoprenoid metabolism as well as stress response pathways. This set included four genes in amino acid biosynthesis, one in phospholipid biosynthesis, and one in the stringent response that, to our knowledge, have not previously been implicated in lycopene production. In contrast, genes for which repression decreased yield were limited to isoprenoid biosynthesis, central carbon metabolism, and stress response pathways. Together, our work reveals genetic targets for increasing lycopene yield throughout metabolism, and defines a tractable, generalizable approach to mapping genetic factors that modulate biosynthetic pathway yield.

## 1. Introduction

Isoprenoids are a large and diverse class of natural products with value as therapeutics, nutraceuticals, cosmetic additives, fragrances, and biofuels (Imran et al., 2020; Kong et al., 2010; Pérez-Gálvez et al., 2020). The abundance of isoprenoid and terpenoid compounds in natural sources is typically low, and in many cases, the extraction process can be labor-intensive and/or toxic (Song et al., 2016). These challenges have positioned microbial biosynthesis of isoprenoids, particularly in the well-characterized microbial host *Escherichia coli*, as an attractive and sustainable alternative. *E. coli* natively synthesizes the universal C5 isoprenoid precursors isopentenyl diphosphate (IPP) and dimethylallyl diphosphate (DMAPP) through the methylerythritol phosphate (MEP) pathway, which converts glyceraldehyde-3-phosphate (G3P) and pyruvate into IPP/DMAPP (Zu et al., 2020) (**Figure 1A**). Introduction of heterologous genes into *E. coli* enables conversion of these molecules into high-value compounds like the taxol precursor taxadiene (Ajikumar et al., 2010), a diversity of carotenoids (Schmidt-Dannert et al., 2000; Umeno et al., 2005), and the potential D2 diesel precursor bisabolene (Alonso-Gutierrez et al., 2018).

**Fig. 1.**
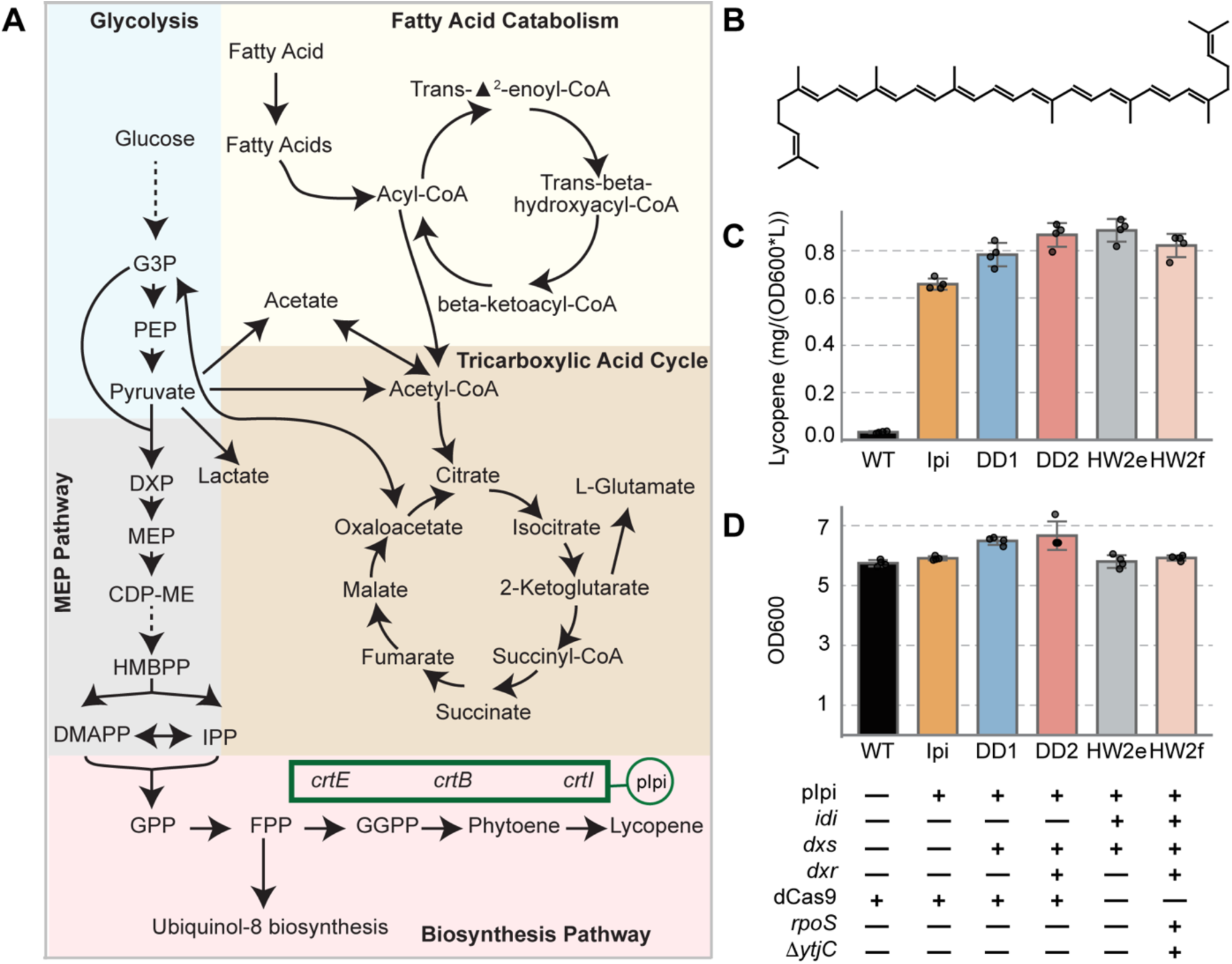
Strain Engineering and Lycopene Pathway Optimization. (**A**) *E. coli* central carbon metabolism and lycopene production. The final five steps of lycopene biosynthesis are indicated with pink shading. All reactions are native to *E. coli* strain MG1655, except *crtEBI* which are provided on the pAC-LYCipi plasmid (abbreviated pIpi above and indicated with a green box). (**B**) Chemical structure of lycopene, a 40-carbon aliphatic hydrocarbon with 11 conjugated double bonds. (**C**) Lycopene production in the strains developed for this study (DD1, DD2) compared to *E. coli* MG1655 (WT), MG1655 transformed with the plasmid encoding *crtEBI* (Ipi), and two strains with selectively modified ribosome binding sites optimized to increase lycopene yield (HW2e, HW2f). Bar height corresponds to mean lycopene yield after 48 hours of culture across four replicates (lycopene mg normalized by volume and OD_600_). Replicates are shown as individual dots; error bars span one standard deviation. (**D**) Optical density at 600 nm following 48 hours of culture in LB media. Again, bar height corresponds to the average across four replicates (replicates are shown as individual dots, error bars span one standard deviation). The table below indicates the presence or absence of engineered genetic elements intended to enhance lycopene production (pIpi = pAC-LYCipi plasmid. dCas9 = genomically integrated, tetracycline inducible dCas9. Δ*ytjC* = elimination of *ytjC* gene function. Rows *idi*, *dxs*, *dxr*, and *rpoS* all indicate the presence/absence of an enhanced translation efficiency ribosome binding site (RBS) upstream of the respective gene.)

However, biosynthesis of many isoprenoids is carbon intensive. Prenyl diphosphate synthases condense 5-carbon IPP and DMAPP units to form linear and often long prenyl chains, which are then cyclized and/or otherwise modified to yield diverse isoprenoids. This consumption of 5-carbon units can introduce significant metabolic burden on *E. coli* central carbon metabolism (Farmer and Liao, 2001; Vadali et al., 2005). Here we examine genetic factors that enhance isoprenoid production by taking lycopene as a convenient and quantifiable colorimetric indicator of IPP/DMAPP biosynthesis. Lycopene is a 40-carbon isoprenoid with a deep red color owing to its conjugated double-bond system (**Figure 1B**). In endogenous microbe (e.g. *Pantoea agglomerans*) and plant producers, lycopene is primarily an intermediate metabolite in the synthesis of other carotenoids and phytohormones, and functions as a photo-protective antioxidant (Cazzonelli and Pogson, 2010; Umehara et al., 2008). Industrially, it is widely used as a food colorant and cosmetic additive, yet extraction from natural sources is inefficient, and large-scale chemical synthesis is both toxic and solvent-intensive. Heterologous expression of the downstream genes *crtE*, *crtB*, and *crtI* from *Pantoea agglomerans* in *E. coli* enables lycopene biosynthesis through condensation of IPP/DMAPP into geranylgeranyl pyrophosphate (GGPP) and its desaturation into lycopene (Cunningham et al., 2007a) (**Figure 1A**). Like other isoprenoids, this process diverts substantial metabolic flux from central metabolism, and competition for pyruvate, glyceraldehyde-3-phosphate (G3P), and acetyl-CoA often limits production efficiency. Moreover, excess accumulation of IPP and prenyl diphosphate intermediates can exert cytotoxic effects, impairing growth and reducing productivity (Ajikumar et al., 2010; Alper et al., 2005; George et al., 2018).

Changes in both gene expression and environment can help address these challenges by reducing metabolic burden and improving precursor availability. For example, redistributing carbon flux to increase G3P availability relative to pyruvate significantly enhanced lycopene production in *E. coli* (Farmer and Liao, 2001). Likewise, disrupting competing pathways at the pyruvate and acetyl-CoA nodes (*ackA-pta*, *nuo* mutants) increased lycopene production (Vadali et al., 2005). Additionally, using auxiliary carbon sources such as free fatty acids or waste oils boosted lycopene production in appropriately engineered cells (Liu et al., 2020). Together, these findings illustrate that isoprenoid biosynthesis is broadly coupled to central carbon metabolism, including the tricarboxylic acid (TCA) cycle, glycolysis, and fatty acid catabolism (**Figure 1A**). Consequently, optimizing isoprenoid production requires going beyond localized, rationally-directed tuning of expression for specific genes in the MEP pathway and *crt* operon to comprehensively explore the connection between metabolism and pathway yield.

Recent advances in CRISPR-based functional genomics provide a new route to map the relationship between metabolism and isoprenoid yield at scale. CRISPR interference (CRISPRi) enables programmable repression of endogenous genes without altering the genomic sequence. In this system, the 20-nucleotide guide region of a single guide RNA (sgRNA) is used to target a catalytically dead Cas9 endonuclease (dCas9) to the genome, where it binds and inhibits transcription (Hawkins et al., 2015; Qi et al., 2013). Coupling CRISPRi with next-generation sequencing (NGS) makes it possible to measure the growth rate effect of thousands of repressions in a single experiment, an approach sometimes called CRISPRi-seq. Because this approach confers inducible repression, it can be used to quantify the functional consequences of perturbing essential genes, unlike transposon-based screens (Alper and Stephanopoulos, 2008). Moreover, introducing mismatches into the sgRNA enables stepwise modulation of gene expression, making it possible to quantitatively examine the effects of gradated expression changes for balancing growth and metabolite production (Hawkins et al., 2020; Mathis et al., 2021; Otto et al., 2024; Vigouroux et al., 2018). CRISPRi-seq has now been used to conduct large scale screens of the impact of gene repression on growth, morphology, and antibiotic resistance (Hawkins et al., 2020; Koo et al., 2024; Peters et al., 2016; Silvis et al., 2021). However, it has found more limited application in biosynthetic pathway optimization. Prior work has used CRISPRi repression of a small number of genes to change lycopene flux in *E. coli* as a proof-of-concept for new sgRNA designs (Byun et al., 2023). Another study performed a CRISPRi screen targeting 74 genes to identify modulators of native carotenoid production in *Corynebacterium glutamicum* (Göttl et al., 2021). However, these workflows did not integrate a NGS step to comprehensively map sequence/yield relationships, limiting library size and throughput.

Here we conduct an expanded CRISPRi screen across 180 *E. coli* genes and establish a 96 well plate-based assay paired with NGS for improved throughput. We observed that the phenotypic response to gene repression was strongly dependent on the timing of CRISPRi induction. Consistent with prior work, we found a tradeoff between lycopene yield and biomass production (Farmer and Liao, 2001; Sun et al., 2014; Wang et al., 2020). We identified 31 genes that significantly modify lycopene production, including genes in central carbon metabolism, branched chain amino acid biosynthesis, isoprenoid biosynthesis, fatty acid biosynthesis, and both the stringent and SOS response pathways. Our work defines a new set of metabolic targets for optimizing lycopene (and isoprenoid) biosynthesis and opens the door to using high-throughput machine learning strategies to discover the relationship between gene expression variation and biosynthetic pathway yield.

## 2. Materials and methods

### 2.1 Bacterial strains and Media

All cloning and library preparation was performed in *E. coli* XL1-Blue cells *(recA1 endA1 gyrA96 thi-1 hsdR17 supE44 relA1 lac [F’ proAB lacIq ZΔM15 Tn10 (Tetr)])*. All CRISPRi experiments were carried out in *E. coli* K-12 MG1655 cells (F- ilvG- rfb-50 rph-1 HK022 attB::dCas9). For indicated experiments, we also used isogenic MG1665::dCas9 derivatives with engineered ribosome-binding site (RBS) substitutions: one strain with an updated RBS upstream of *dxs* (MG1655 HK022 attB::dCas9 RBS^*dxs*, Fig. S1B), and a second strain with updated RBS upstream of *dxs* and *dxr* (MG1655 HK022 attB::dCas9 RBS^*dxs*,*dxr*: Fig. S1C). We refer to these strains as DD1 and DD2 respectively, details of strain construction follow in section 2.2. All edited loci were sequence-verified. A complete list of cloning and sequencing primers can be found in **Table S1**.

### 2.2 Genomic integration of high-efficiency RBS in the MEP pathway

We used Oligonucleotide Recombineering followed by Bxb-1 Integrase Targeting (ORBIT) to modify the RBS for the MEP pathway genes *dxs* and *dxr*. This process for genomic engineering requires three components: a helper plasmid (pHelper), an integrating plasmid (pInt), and a targeting oligo. The targeting oligo encodes a 38-bp attB site flanked by homology arms, which is incorporated into the genome during replication by the single-stranded DNA annealing protein CspRecT. The plasmid pInt carries the cognate attP site, which recombines with the oligo via the Bxb1 recombinase, resulting in stable genomic integration. Expression of CspRecT and Bxb1 is independently induced by *m*-toluic acid and L-arabinose, respectively from pHelper. The pHelper plasmid has a temperature sensitive origin of replication, and can later be cured from the cells by growth under a non-permissive temperature (37–42 °C, Saunders and Ahmed, 2024).

To achieve scarless editing of the RBS sequence with ORBIT, we followed a two-step editing process. In the first step, we used an attB-containing targeting oligo to replace the endogenous RBS with pInt carrying both a selectable antibiotic marker (KanR) and the counter-selectable *sacB* gene. The *sacB* gene confers sucrose sensitivity, such that cells retaining the marker grow more slowly in the presence of sucrose. In the second step of editing, we replaced the genomically integrated pInt with a clean targeting oligo lacking an attB site and encoding the new RBS. These cells were then grown on sucrose to select for colonies that underwent successful excision of pInt through a second-round ORBIT event with the clean attB-free oligo (Saunders and Ahmed, 2024).

#### 2.2.1 ORBIT targeting oligo Design

Briefly, for ORBIT in *E. coli* K-12 MG1655, targeting oligos must anneal to the lagging strand template, which is determined by replichore orientation. Replichore 1 (<1.59 Mb or >3.92 Mb) requires “–“ strand homology, while replichore 2 (1.59–3.92 Mb) requires “+” strand homology. The orientation of the attB site is also important and is typically aligned with the target gene. To simplify this process, a dedicated web application (<u>ORBIT oligo design tool</u>) generates oligos that satisfy these strand and orientation rules. Users can select genes of interest in a genome browser, adjust homology arm length or attB direction, and directly copy the resulting 5′–3′ sequence for ordering. We used this tool to design targeting oligos for the *dxs* and *dxr* loci to incorporate the optimized RBS sequence. For each scarless genomic edit, two oligos were prepared. Oligo 1 (120 nt) contained a 38-bp attB site flanked by optimized RBS sequence and homology arms to the target locus. Oligo 2 (90–120 nt) contained the optimized RBS sequence and appropriate homology arms, but no attB site (**Table S1, Figure S1B,C**).

#### 2.2.2 ORBIT-induced electrocompetent cells preparation, transformation, and recovery

Targeting oligo 1 (1µM) and the integrating plasmid (pINT_attP1_sacB_kanR, 100 ng) were co-transformed into freshly prepared m-toluic acid-induced MG1655::dCas9 competent cells harboring the temperature sensitive helper plasmid pHelper_TS_V2_AmpR as described (Saunders and Ahmed, 2024). Cultures were incubated at 30 °C while shaking for 1 h prior to plating on LB agar plates with kanamycin and ampicillin selection and grown at 30 °C overnight. Successful first-round ORBIT integrations of the pINT plasmid were confirmed by colony PCR using locus-specific primers for either *dxs* or *dxr* (Table S1). These colonies, which still carried the pHelper_TS_V2 plasmid, were then used for a second round of ORBIT.

These colonies were subsequently used to prepare ORBIT-induced electrocompetent cells as described above, following the same induction and transformation procedures, with growth maintained in LB supplemented with both kanamycin and ampicillin. After transformation and electroporation with the clean attB-free Oligo 2, cells were recovered under the same conditions but were now plated on LB agar containing 10% sucrose for counter-selection against pINT integration. Again, colony PCR was performed to detect the scarless edit, yielding an amplicon of ∼300–400 bp rather than the ∼4 kb band observed after the first-round integration. Following PCR verification, the temperature-sensitive helper plasmid (pHelper_TS_V2_ampR) was cured by growing colonies at non-permissive temperatures (37–42 °C Saunders and Ahmed, 2024). Colonies were subsequently streaked in duplicate on LB agar with and without ampicillin at 30 °C to screen for loss of the plasmid. The RBS of *dxs* was edited first to create strain DD1. The strain DD1 was then further edited to modify the RBS of *dxr* and create DD2. Colonies of both DD1 and DD2 were further validated by Sanger sequencing of the RBS loci and whole- genome sequencing to confirm successful genome editing and to rule out off-target effects in DD1 and DD2 strains (Figure S1).

### 2.3 Whole-genome DNA extraction, sequencing, and *breseq* analysis

Genomic DNA was extracted from the *dxs* and *dxr* strains using the Quick-DNA Fungal/Bacterial Miniprep Kit (Zymo Research, D6005). Whole-genome sequencing was performed by Plasmidsaurus (Eugene, OR, USA), and raw reads were provided in FASTQ format. Sequence analysis was carried out using *breseq* (Barrick Lab; https://barricklab.org/twiki/bin/view/Lab/ToolsBacterialGenomeResequencing) within the UTSW BioHPC computing environment. FASTQ files were first placed in the designated working directory, and *breseq* was executed using two computational cores with the following command: “breseq -j 2 -x -r MG1655_dCas9_reference_genome_circular.gb sample.fastq.gz” where the reference genome was the circularized MG1655–dCas9 sequence (downloaded in GenBank format) and the input FASTQ files corresponded to the Oxford Nanopore sequencing reads (Plasmidsaurus). Analysis runs were typically completed within ∼1 h. Output files were subsequently inspected to identify mutations, confirm targeted modifications, and rule out off-target effects.

### 2.4 High-throughput CRISPRi Induction

The DD1 or DD2 strain was transformed with CRISPRi sgRNAs encoded on the pCRISPR3 plasmid by electroporation. Transformants were grown on LB agar plates containing kanamycin and chloramphenicol to select for pCRISPR3 and pAC-LYCipi respectively. Individual colonies were picked and grown overnight in 96-deep-well plates (Fisher Scientific, AB-0932) containing 1 mL LB medium supplemented with the same antibiotics at 30 °C on a microplate shaker (Fisher Scientific, 13687708). From each overnight culture, 15 µL inoculum was transferred into fresh deep-well plates containing 1 mL LB with kanamycin and chloramphenicol and incubated for 2 h at 30 °C to allow a return to logarithmic growth phase. Following outgrowth, optical density at 600 nm (OD₆₀₀) was measured for each culture in a flat-bottom 96-well plate using an Agilent BioTek multimode microplate reader, and cultures were adjusted to achieve a starting OD₆₀₀ of 0.005 in new deep-well plates containing 1 mL LB with kanamycin and chloramphenicol. Cultures were grown for 4 h at 30 °C prior to dCas9 induction with anhydrotetracycline at a final concentration of 50 ng/mL; cultures were then incubated for a total of 48 h at 30 °C. Final OD₆₀₀ was measured, and lycopene content was quantified by acetone extraction followed by absorbance at 475 nm. For induction time-course experiments (**Figure 2**), cultures were seeded at OD₆₀₀ = 0.005 as above, and CRISPRi was induced at 0, 4, 8, or 24 h. Cultures were then allowed to grow for 48, 44, 40, or 24 h, respectively, such that the total outgrowth period for each condition was 48 h.

**Figure 2.**
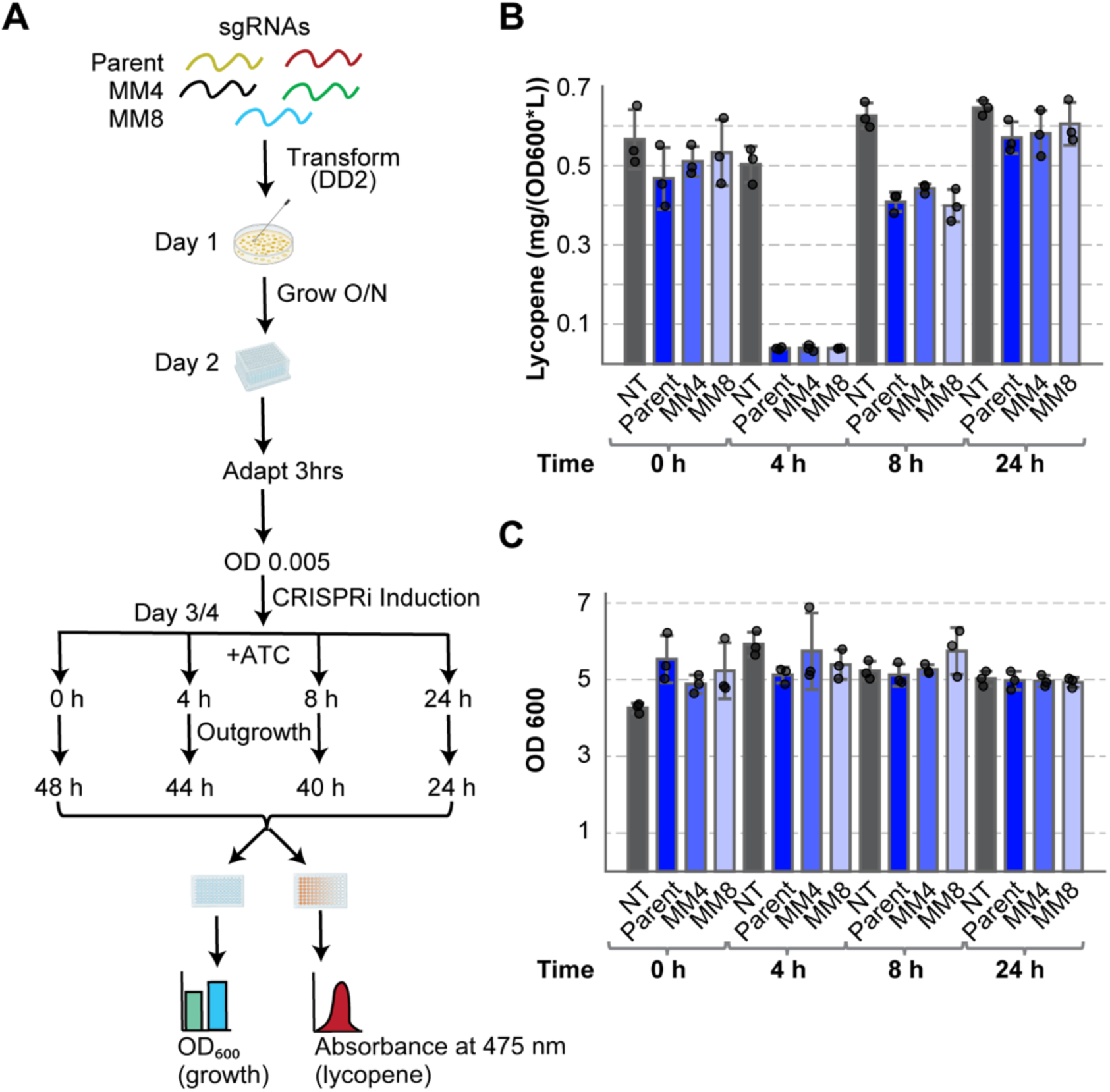
Optimizing CRISPRi Induction Timing for Strong Repression Effects. (**A**) Experimental workflow used to evaluate the effect of CRISPRi induction timing on lycopene production. The DD2 *E. coli* strain was transformed with sgRNAs targeting *dxs*, including a parent guide and mismatch variants (MM4 and MM8) designed to produce gradated levels of repression. Cultures were grown overnight, back-diluted and allowed to re-enter log phase growth (3 hour adaptation), then diluted to OD₆₀₀ = 0.005 in fresh media. CRISPRi was induced with anhydrotetracycline (+ATC) at the indicated timepoints (0, 4, 8, or 24 h) during outgrowth. Cells were harvested after a total of 48 h growth for measurement of biomass (OD_600_) and lycopene production (absorbance at 475 nm). (**B**) Lycopene yield normalized to OD_600_ (mg / (L * OD_600_)) for cells expressing the non-targeting control (NT), the parent *dxs*-targeting sgRNA, or mismatch variants (MM4, MM8) at different CRISPRi induction timepoints. Bars show mean values (n = 3 biological replicates) with individual measurements as black dots. (**C**) Biomass production (OD_600_) measured for the same conditions shown in panel B. Induction of CRISPRi after 4 h of outgrowth produced the largest change in lycopene yield while maintaining robust cell growth. Bars show mean values (n = 3 biological replicates) with individual measurements as black dots.

### 2.5 Lycopene extraction and quantification

Cultures (650 µL in size) were transferred to Eppendorf tubes and centrifuged for 5 min at 21k RCF, washed once with PBS, and resuspended in 650 µL acetone. Samples were vortexed, incubated at 55 °C in a water bath for 15 min, cooled, and centrifuged for 7 min (21k RCF). The resulting supernatant was transferred to fresh Eppendorf tubes, and 200 µL was transferred to a quartz 96-well plate to measure absorbance at 475 nm (Jung et al., 2016). Lycopene standard (Sigma-Aldrich, USA) was used for calibration, and all steps were performed under dark conditions to prevent light-induced isomerization. Lycopene yield is commonly reported as a fraction of the dry cell weight (Jung et al., 2016; Li et al., 2024; Wu et al., 2018), culture volume (Farmer and Liao, 2001; Ren et al., 2023; Vadali et al., 2005), or both (Liu et al., 2020; Xu et al., 2018; Zhan et al., 2024). Following from a previous study (Yuan et al., 2006), we quantified lycopene yield as mg/(L* OD_600_) with the goal of accounting for perturbations in biomass while avoiding the more laborious process needed for dry cell weight.

### 2.6 CRISPRi sgRNA library design

We followed established criteria for mismatch-CRISPRi in *E. coli* to design the gene-specific homology region of each sgRNA in our library. More specifically, sgRNAs targeted the non-template strand adjacent to a CCN PAM sequence, had 45–80% GC content, and excluded poly-T tracts greater than 4 nucleotides. All candidate sgRNA sequences were screened against the MG1655 genome by BLASTn, and those with ≥75% identity at a PAM-adjacent off-target site were rejected (Mathis et al., 2021; Otto et al., 2024). For CRISPRi timing and protocol optimization experiments, guides were designed against *dxs* using the parent sgRNA closest to the transcription start site along with mismatch variants over the first 4 and 8 positions (distal to the PAM sequence). Two broader libraries were then constructed. In the small library, *ldhA, ispA, dxs,* and *dxr* were each targeted with three parent sgRNAs with two proximal to the transcription start site and one further away. Seven different mismatch variants were generated per parent by introducing compounding mutations (over the first 2, 4, 6, 8, 10, 12, and 14 positions), yielding 96 guides plus a non-targeting control. In the large library, we initially selected 184 metabolic and regulatory genes; however, four genes were excluded due to design limitations (e.g., absence of unique PAM-proximal sites or sequence similarity to paralogs), yielding a final set of 180 genes. For each gene we designed one sgRNA positioned closest to the start codon and a corresponding mismatch-7 variant, except for *gdhA*, for which the automated design pipeline did not generate a mismatch-7 guide, resulting in a library containing 359 guides in addition to the non-targeting control. The sgRNA libraries were ordered as oligonucleotide pools from IDT, with each oligo containing promoter, sgRNA, and terminator elements.

### 2.7 sgRNA library construction and assembly

Golden Gate cloning was used to assemble sgRNA libraries from oligonucleotide pools into the pCRISPR3 barcoded destination vector. BsaI overhangs were created by using the GG_univ_F and GG_univ_R primers (**Table S1**). Two microliters of each purified Golden Gate reaction was transformed into 100 µL XL1-Blue chemically competent *E. coli*. Following a 1-hour recovery in SOB medium at 37 °C with shaking at 220 rpm, a 1:10 and 1:100 dilution of the transformation was plated on LB-kanamycin agar plates to estimate cloning efficiency. The remainder of the transformation was expanded overnight in SOB medium supplemented with 35 µg mL⁻¹ kanamycin, after which glycerol stocks were prepared for long-term storage. A subset of colonies was submitted for Sanger sequencing to confirm correct incorporation of sgRNA sequences from the oligo pool. Library coverage was calculated relative to no-insert golden gate controls and consistently exceeded tenfold representation of the theoretical library size, confirming robust diversity across constructs. The extracted plasmid DNA was subsequently transformed into the DXR-optimized DD2 *E. coli* strain carrying the pAC-LYCipi plasmid (**Figure S1A**). Single colonies were isolated for downstream CRISPRi experiments and MiSeq analysis to identify the sgRNA.

### 2.8 Amplicon sample preparation for NGS

Following CRISPRi induction and outgrowth in 96-deep-well plates (see section 2.4), DNA amplicons for Illumina sequencing were prepared as described previously (Mathis et al., 2021; Otto et al., 2024). Briefly, DNA was extracted by boiling cell pellets in sterile water at 95 °C for 5 minutes, and clarified lysates were then used directly as templates for PCR to add the Illumina adaptors. In the first amplification step, universal TruSeq adapter sequences were appended to the sgRNA cassette using Q5 Hot Start High-Fidelity DNA Polymerase (NEB, M0493) with adapter-specific primers. For plate-level identification in the large-scale genomic screen, an additional 8 bp barcode was incorporated into the TruSeq forward primer (Table S1), enabling unambiguous assignment of sequencing reads to their originating plate. The TruSeq universal reverse primer was used without modification. The TruSeq-adapted products were then subjected to a second PCR to append unique i5 and i7 index sequences, which encoded the well position on the 96-well plate. Following amplification, double-stranded DNA concentrations were determined using both PicoGreen (Thermo Fisher, P7581) and Qubit dsDNA HS assays (Thermo Fisher, Q32851). For PicoGreen, lambda DNA standards (2 ng µL⁻¹ in TE buffer) were used to generate calibration curves in black-walled, clear-bottom 96-well plates. DNA samples (1 µL diluted in 99 µL TE) were incubated with 100 µL PicoGreen reagent (1:200 in TE), protected from light, and fluorescence was read on a multimode plate reader. In parallel, Qubit assays were performed with three dilutions (1:1, 1:10, and 1:100) of each sample by mixing 10 µL DNA with 190 µL Qubit working reagent and incubating for 2 min in the dark. Measured concentrations were adjusted by the dilution factor and compared with PicoGreen values to ensure accuracy. Based on these quantifications, Round 2 PCR products were pooled to yield equimolar contributions of ∼1 µg dsDNA per library. Pooled libraries were concentrated by gel extraction, purified with a Zymo DNA Clean & Concentrator kit, and quality was confirmed by Nanodrop A260/280 ratio prior to sequencing. Final library concentrations were normalized to ∼2 nM and denatured with NaOH prior to loading. PhiX control DNA (Illumina) was spiked at 20% to provide sequence diversity. Sequencing was carried out on an Illumina MiSeq using the TruSeq DNA PCR-free kit with 2 × 150 bp paired-end reads. Libraries were prepared at 8 pM loading concentration in hybridization buffer, and run following the standard MiSeq workflow.

### 2.9 sgRNA assignment by sequencing

#### 2.9.1 Read mapping and quality filtering

Raw FASTQ files resulting from Illumina sequencing were parsed into paired read 1 (R1) and read 2 (R2) sets for each well. For each read pair, sgRNA sequences were identified using regex-based searches anchored to invariant flanking regions of the sgRNA cassette. To ensure robust assignment, a maximum Hamming distance of three mismatches from the expected sgRNA window was permitted. Sequence quality filtering was applied at the base level, requiring all bases at the intended mismatch site and its ±1 neighboring nucleotides to have a Phred quality score ≥30. Reads passing this threshold were retained for downstream analysis.

Assignment of reads to individual sgRNAs was achieved by mapping the 20-nt guide sequence against a reference sgRNA library. To minimize spurious calls, only sgRNAs supported by ≥100 reads and accounting for ≥95% of total reads in a well were accepted. Wells containing multiple high-frequency sgRNAs were flagged as potential cross-contamination and excluded. Filtered counts were summarized into per-sample tables, with both raw and processed outputs exported in .csv and Excel formats for reproducibility.

#### 2.9.2 Differences between small- and large-library pipelines

- *Small-library experiments*: Wells were tracked without explicit plate-level barcoding. Assignment required the detection of a valid 20-nt sgRNA homology region paired with the TruSeq i5/i7 index pairs to a well.
- *Large-library experiments*: In addition to the above, an 8-bp plate-level barcode was incorporated into the TruSeq forward primer. Four distinct barcodes (AATAGGCG, AATGAGCC, AATCTCCG, AAGAAGGC) were used to distinguish the four 96-well plates in our experiment (designated F1–F4). These barcodes were parsed from R1 reads and used to unambiguously assign samples to their originating plate. The remainder of the workflow mirrored that of the small library, with sgRNA sequences identified from R2. Per-plate read distributions were analyzed to confirm balanced representation across wells and plates.

Together, these criteria ensured accurate and reproducible assignment of sgRNA identities across hundreds of wells. By enforcing Hamming distance tolerance, stringent base quality thresholds, and read-count filters, the pipeline minimized misidentification due to sequencing errors. The inclusion of plate-level barcoding in the large-library design further enabled robust deconvolution of multiplexed samples, thereby supporting high-throughput CRISPRi library screening at scale.

## 3. RESULTS

### 3.1 A lycopene-producing *E. coli* chassis strain for CRISPRi screening

We sought a relatively high-yield chassis strain as a starting point for CRISPRi screening under the reasoning that this would facilitate detection of small changes in lycopene production. To achieve this, we began with a MG1655 K12 *E. coli* strain, containing a genomically integrated dCas9 enzyme under control of a tetracycline-inducible promoter (Mathis et al., 2021; Vigouroux et al., 2018). Prior work indicated that K12 *E. coli* are a favorable strain for lycopene production (Ren et al., 2023). We transformed this strain with the plasmid pAC-LYCipi (**Figure S1A**), encoding lycopene synthesis genes *crtEBI* and *idi* from *Pantoea agglomerans*, to yield strain *Ipi* (Cunningham et al., 2007b). This strain contained the minimal genetic components required for lycopene synthesis and produced detectable lycopene and grew to the same liquid culture optical density as the wild type (**Figure 1C**,**D**). To further enhance lycopene yield, we took inspiration from prior directed evolution work where Wang and co-workers found that increasing the translation efficiency of the ribosome binding sites (RBS) for the MEP pathway genes, *dxs* and *dxr,* enhanced lycopene yield (Wang et al., 2009). Using a genome engineering approach called ORBIT (Saunders and Ahmed, 2024), we replaced the native RBS of *dxs* with a sequence predicted to increase translation efficiency ∼5000-fold (**Figure 1C, Figure S1B**) (Salis, 2011), generating strain DD1. A second ORBIT edit replaced the native RBS of *dxr* with an optimized RBS (∼6000-fold) to generate strain DD2 (**Figure 1C,D**, **Figure S1C**). Under our small-scale culture conditions, DD2 produced lycopene at levels comparable to previously reported laboratory-evolved strains HW2e and HW2f (**Figure 1C**) (Wang et al., 2009). These results establish DD2 as a high-yield chassis suitable for CRISPRi screening.

### 3.2. CRISPRi induction timing impacts the effect of gene repression on lycopene production

Next, we developed our CRISPRi screening protocol in the context of strain DD2. A key element is to establish the timing of CRISPRi induction relative to subsequent culture outgrowth and measurement of lycopene yields. Early induction of CRISPRi repression for essential genes may result in lethality, and/or strong selection for CRISPRi “escapers” — variants that evade repression by mutating the dCas9 element, the sgRNA, or the genomic target sequence (Mathis et al., 2021). In contrast, late induction of CRISPRi might achieve only limited upregulation of dCas9 production and/or miss the opportunity to shift key metabolites away from biomass production and towards lycopene biosynthesis during exponential growth. Under our experimental conditions, lycopene production becomes most apparent after the cells have reached stationary phase, meaning that the experimental readout may only become apparent many hours after the CRISPRi perturbation.

To define and optimize our CRISPRi protocol, we targeted repression of *dxs* under the expectation that *dxs* repression would yield a strong decrease in lycopene yield. We designed sgRNAs with a 20 bp homology region complementary to *dxs*. We also introduced four and eight sequential bp mismatches distal from the PAM-adjacent end of the homology region to achieve attenuated CRISPRi activity against *dxs* (Mathis et al., 2021). Plasmids expressing these sgRNAs, along with a non-targeting control (NT), were introduced into DD2. Cultures were induced with anhydrotetracycline at 0, 4, 8, or 24 hours following back-dilution and low-density inoculation into deep well culture plates (**Figure 2A**). We harvested cells after 48 hours of total growth, and measured biomass (OD₆₀₀) and lycopene production.

We observed the strongest reduction in lycopene yield when CRISPRi was induced after four hours of growth, followed by induction at eight hours (**Figure 2B**). OD_600_ measurements indicated that these induction points occurred during exponential growth (**Figure S2A**). Despite the strong effect on lycopene production, CRISPRi induction had little effect on final culture density (**Figure 2C**). Given that *dxs* is an essential gene, we presumed that either (1) the cells were able to grow given an incomplete knockdown (rather than knockout) and/or (2) mutations emerged that disrupted CRISPRi repression and enabled growth as the experiment proceeded (so-called “escaper” mutations). Similar results were obtained in strain DD1 (**Figure S2B,C**), and so four hours of outgrowth was chosen as our induction timepoint moving forward. Together, these findings demonstrate that CRISPRi induction timing strongly influences screening outcomes.

### 3.3 A high throughput sequencing-based CRISPRi Screen for lycopene production

With the goal of expanding our screen to hundreds of genes across metabolism, we coupled our CRISPRi selection and lycopene measurements to NGS for sgRNA identification. Conceptually, our approach is similar to the every variant sequencing (evSeq) strategy introduced by the Arnold lab (Wittmann et al., 2022). The basic idea is to first generate barcoded PCR amplicons spanning the sgRNA homology region, and where each barcode pair uniquely maps to a specific well of the 96-well plate. Then, low-cost short-read NGS of the pooled amplicons is used to identify the sgRNA associated with each well (**Figure 3A**). This process allows high-throughput, cost-effective mapping of every sgRNA repression to lycopene yield phenotype.

**Figure 3.**
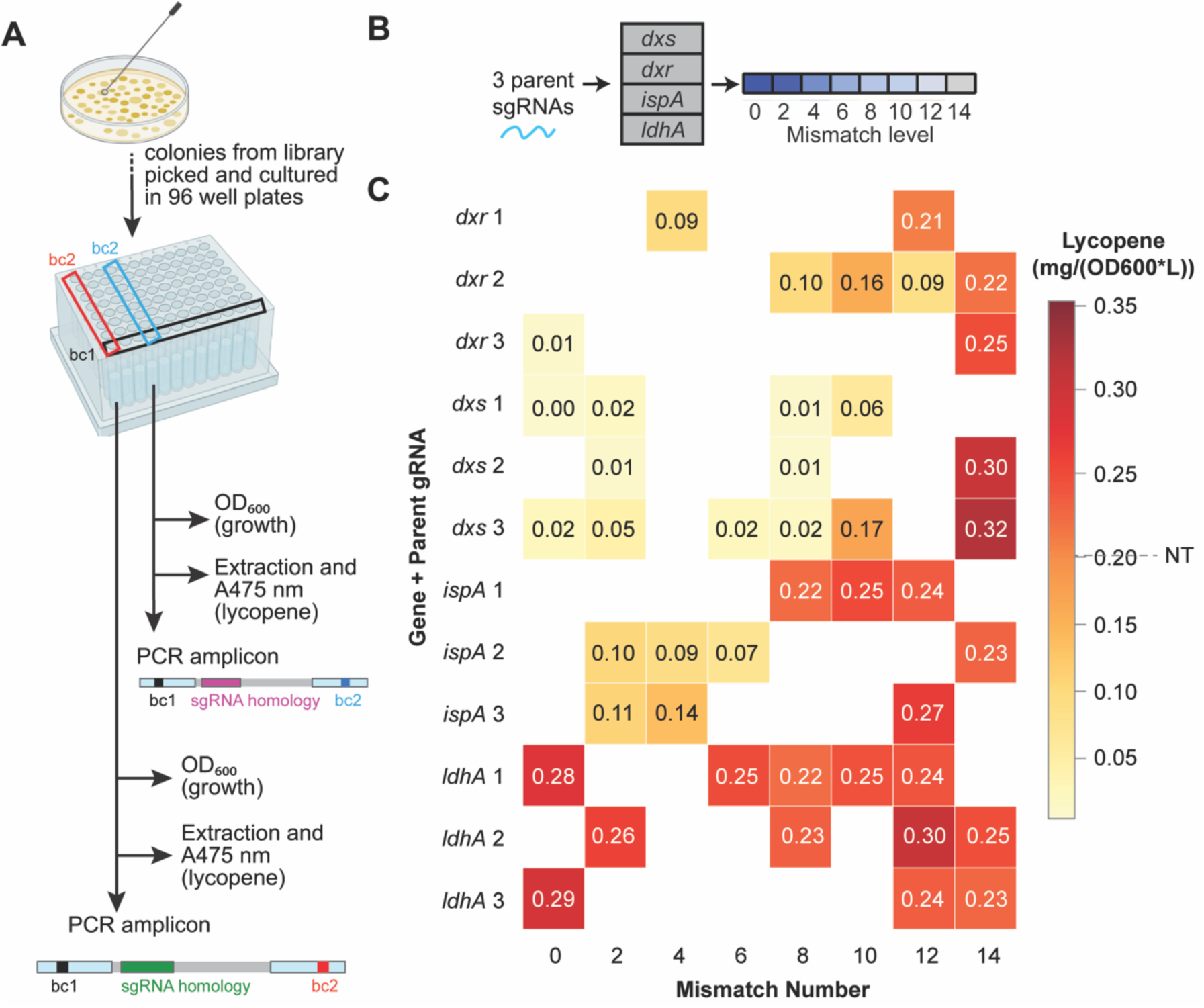
Establishing 96 well plate mismatch CRISPRi Screen using four target genes. (**A**) Experimental workflow for validating mismatch-CRISPRi modulation of lycopene production. Strains expressing sgRNAs were grown in microplates, and growth (OD_600_ and lycopene levels were measured after 48 h by extraction and absorbance at 475 nm. sgRNA identities were mapped by PCR amplification and NGS. DNA barcodes indicated the column (bc2) and row (bc1) on the plate. (**B**) Design of the mismatch sgRNA library. For each gene (*dxs, dxr, ispA,* and *ldhA*), three parent sgRNAs were used to generate variants containing 0–14 mismatches relative to the target sequence. (**C**) Heatmap of lycopene production in strain DD2 across sgRNA variants targeting *dxs, dxr, ispA,* and *ldhA*. Rows indicate parent sgRNAs (grouped by gene), and columns indicate mismatch number. Values represent mean lycopene production (mg L⁻¹/ OD_600_) from 1–4 biological replicates. The dashed line on the scale indicates lycopene yield for the non-targeting (NT) reference strain.

To establish this approach, we constructed a focused sgRNA library targeting four metabolic genes expected to have varied effects on lycopene yield: *dxs*, *dxr*, *ispA*, and *ldhA*. The first three *dxs*, *dxr*, and *ispA* genes encode metabolic enzymes that make precursors for lycopene production, while *ldhA* diverts pyruvate away from lycopene production. We reasoned that targeting these genes with CRISPRi would test our ability to detect both decreases and increases in lycopene yield upon gene repression. We designed three parent sgRNAs per gene, each targeting a distinct PAM site in the 5’ region of the targeted gene. To test if we could see fine-scale effects by attenuating the repression strength we then added 2, 4, 6, 8, 12, and 14 sequential mismatches on the PAM-distal side of the homology regions for each of these guides (**Figure 3B**) yielding a 96 member sgRNA library (**Table S2**). The nontargeting (NT) sgRNA was separately transformed into DD2 (to create a strain NT) to ensure a reference in the screen. As additional controls, we included strains HW2e and HW2f as high-lycopene benchmarks, and strain DD2 carrying the same pCRISPR3 sgRNA plasmid with full homology targeting *dxs* used in the induction-timing experiment (**Figure 2**), as a CRISPRi repression control. After transformation and plating on selective agar, 92 single colonies were picked and cultured according to the protocol outlined in **Figure 2A**, with CRISPRi induction at 4 h. Following 48 h of growth in deep-well plates, lycopene yield and OD_600_ were measured for each well, and a subset of cells was collected for amplicon generation. Of the 92 colonies picked, one was excluded due to an error during lycopene extraction and quantification. Amplicon sequencing revealed that 26 wells could not be uniquely mapped to a single sgRNA, leaving 65 wells with confidently assigned guides. Inspection of the 26 wells with non-unique mappings indicated the presence of multiple pCRISPR3 constructs within colonies, likely resulting from excess plasmid during transformation. This issue was corrected in subsequent experiments by reducing plasmid input. The remaining 65 wells mapped to 43 unique guides (1-4wells/guide, mean 1.48), thus 44.8% of the 96-member input library was recovered and measured (**Figure 3C**).

Consistent with our expectation, we observed the lowest lycopene yields for fully on-target (0 mismatch) or low mismatch sgRNAs targeting *ispA*, *dxr*, and *dxs* (**Figure 3C**). These genes encode enzymes directly involved in isoprenoid biosynthesis; therefore, strong repression by CRISPRi reduces lycopene production. Conversely, we observed the highest lycopene yields for on-target and low mismatch sgRNAs targeting *ldhA*. D-lactate dehydrogenase encoded by (*ldhA*) converts pyruvate into lactate; reducing the abundance of this enzyme is expected to preserve pyruvate for lycopene production. When examining the trend in lycopene yield with increasing mismatches, we found a non-monotonic effect of repressing *dxr* and *dxs.* That is, the non-targeting sgRNA gave a lycopene yield of 0.2 mg/(OD600*L) in this experiment. In comparison, modest repression (corresponding to 14 mismatches) slightly improved lycopene production, while strong repression (fewer than 14 mismatches) was deleterious. This suggested that the engineered RBS sequences for these genes resulted in a higher than optimal level of expression for lycopene biosynthesis. Together, these results demonstrated that our screen can quantitatively detect both increases and decreases in lycopene production, and motivated application of this protocol to a larger gene set.

### 3.4. A CRISPRi screen identifies modulators of lycopene yield across *E. coli* metabolism

Next, we performed a CRISPRi screen to broadly interrogate how central carbon metabolism, isoprenoid precursor allocation, and global regulatory networks influence lycopene production in our engineered *E. coli* chassis strain. To accomplish this, we designed a sgRNA library targeting 180 genes spanning central metabolism and regulatory pathways (**Figure 4A, Table S3**). Lycopene biosynthesis proceeds through the MEP pathway, which draws directly from glyceraldehyde-3-phosphate (G3P), pyruvate, and cellular NADPH pools (**Figure 1A**). Accordingly, we designed sgRNAs targeting genes spanning glucose uptake, early glycolytic steps, branch-point regulators, and non-oxidative and oxidative PPP enzymes. We also covered pyruvate-utilizing and acetyl-CoA-generating enzymes, the pyruvate dehydrogenase complex, the TCA cycle, the glyoxylate shunt, and major fermentation pathways. These genes determine whether pyruvate and acetyl-CoA enter respiration, overflow metabolism, amino acid biosynthesis, or the MEP pathway, making them central nodes for tuning precursor availability for lycopene. We also incorporated sgRNAs targeting genes involved in β-oxidation, fatty acid synthesis, phospholipid and cardiolipin biosynthesis, and acetate formation, all of which represent major competing sinks for acetyl-CoA and reducing power. Their inclusion enabled us to assess whether diverting carbon away from these endogenous lipid and membrane-associated pathways could enhance flux toward isoprenoid precursor pools. To directly interrogate steps in isoprenoid metabolism, we targeted nearly all enzymes in the MEP pathway and associated isomerases, prenyltransferases, and ubiquinone/menaquinone biosynthesis enzymes that share intermediates with lycopene synthesis. Finally, genes associated with the SOS response, stringent response, and other global transcriptional regulators were incorporated to test whether broader cellular regulatory programs influence lycopene production. For each of the 180 genes, we designed two sgRNAs: one fully on-target sgRNA and one with seven mismatches. The exception was for *gdhA,* where the seven mismatch sgRNA exhibited off-target homology and was discarded. This resulted in a final library of 359 gRNAs (see also Methods, **Table S3**). These constructs were transformed into chassis strain DD2, and 384 colonies were picked and inoculated into individual wells of 96-well deep-well plates for characterization following the workflow in **Figure 3A**. In our current work, we examined the contribution of metabolic and regulatory genes to isoprenoid biosynthesis in Luria-Bertani (LB) broth, culture conditions in which amino acids are the main source of carbon (as opposed to simple sugars, glycerol, or fatty acids) (Sezonov et al., 2007). Of the 180 targeted genes, we recovered sgRNAs for 138 (**Table S3**). Notably, sgRNA recovery was not significantly associated with gene essentiality, consistent with the inducible CRISPRi system enabling perturbation of essential genes. For each recovered gene we measured endpoint OD_600_ and lycopene yield in triplicate (**Table S4**).

**Figure 4.**
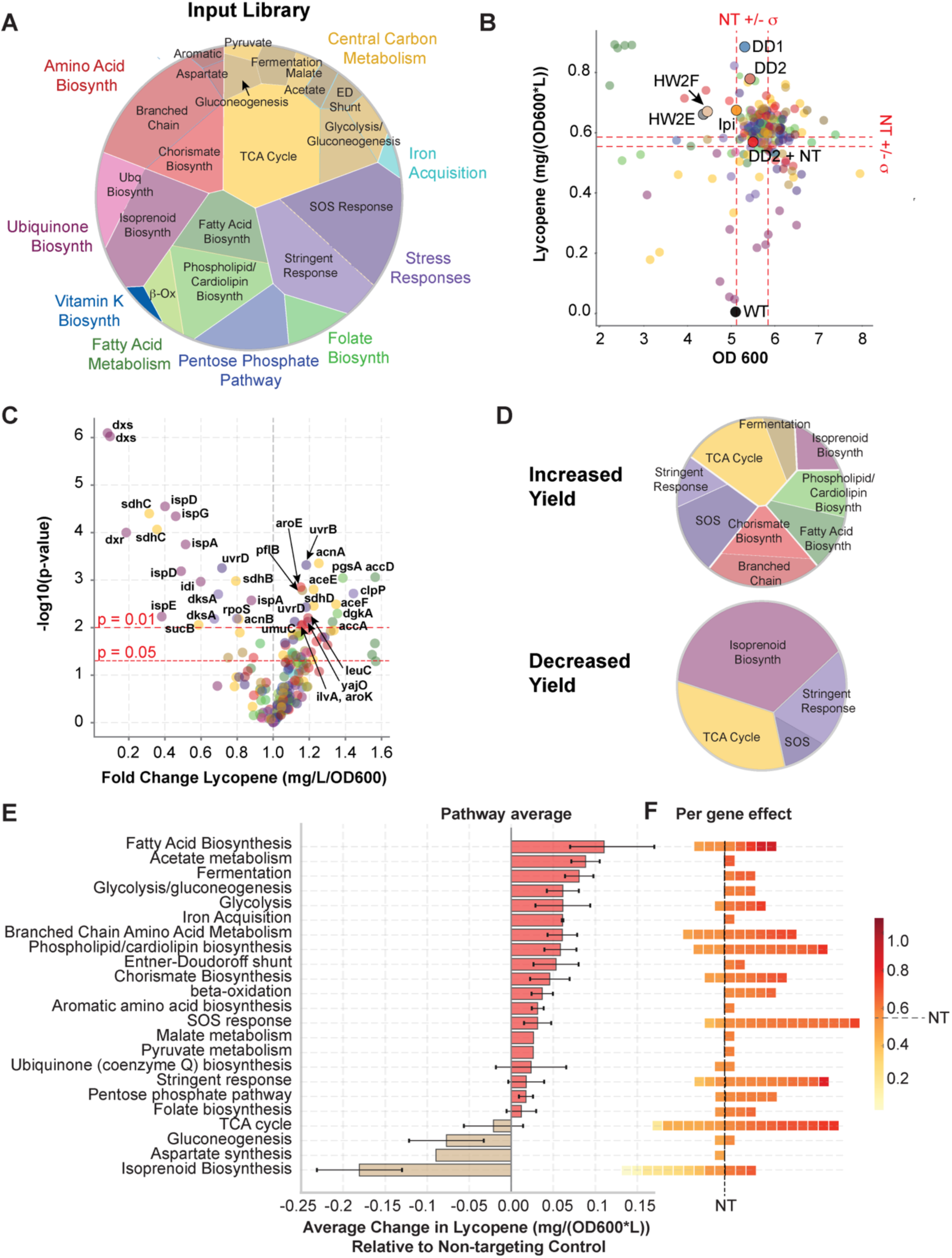
Genome-scale CRISPRi screen identifying metabolic modulators of lycopene production. (A) Composition of the 180 gene CRISPRi sgRNA library, as grouped by metabolic pathway. (B) Cell density (OD_600_) vs lycopene yield (mg/(L*OD_600_)) for all recovered strains. Each point represents a strain expressing a single sgRNA. The non-targeting control strain (NT) is indicated by a red circle, with dashed red lines marking the mean ±1 SD for OD_600_ and lycopene yield. Reference strains (WT, HW2e, HW2f, DD1, and DD2) are indicated by colored circles with black outlines. (**C**) Volcano plot showing fold change in lycopene yield relative to the NT strain (x-axis) and statistical significance (−log10 adjusted p-value, y-axis) from two-sample t-tests with Benjamini–Hochberg correction. Gene names are annotated for all genes with a statistically significant impact on lycopene production. (**D**) Pathway distribution of genes whose repression increased or decreased lycopene yield relative to the NT control. (**E**) Pathway-level analysis showing the mean change in lycopene yield relative to NT, averaged across all genes within each pathway. Error bars indicate standard error. (**F**) Per-gene effects on lycopene yield grouped by pathway. Each square represents the yield associated with the fully on-target parent sgRNA for a gene and is colored according to lycopene production (mg/(OD600*L)). The dashed grey line indicates the lycopene yield of the non-targeting control.

Because high-yield lycopene production in bioreactors will ultimately depend on both pathway yield and overall biomass accumulation, we examined the relationship between the lycopene yield and the endpoint optical/culture density (**Figure 4B**). Most of the screened sgRNAs clustered near the reference strain — the chassis strain DD2 with a non-targeting sgRNA (referred to as NT) — with an endpoint OD_600_ of ∼6 and a lycopene yield slightly above 0.6 mg/(OD_600_*L). Under our culture conditions, NT displayed a slightly lower lycopene yield than DD1 or DD2 alone, suggesting that CRISPRi induction imposes a small metabolic burden. Relative to NT, 92 of the 138 recovered genes had a lycopene yield of at least one standard deviation greater, and 60 genes also displayed at least one standard deviation greater OD_600_. Among these, 38 genes saw increases in both lycopene yield and biomass, representing the most promising candidates for future metabolic engineering.

To identify sgRNAs that significantly altered lycopene production, we performed two-sided t-tests comparing each CRISPRi knockdown strain to the NT chassis strain, followed by Benjamini-Hochberg False-Discovery-Rate correction (**Figure 4C**). We compared these hits to the manually curated annotations of cellular process and metabolic pathway (**Table S3, Figure 4D**). Repression of 13 genes by 19 sgRNAs across 3 cellular processes significantly decreased lycopene yield (**Figure 4D**). Consistent with expectation, many of these sgRNAs targeted isoprenoid biosynthesis genes (10 sgRNAs targeting six genes). The remaining seven genes associated with decreased lycopene production included four TCA cycle genes (*sdhB*, *sdhC*, *sucB*, and *acnB*) and three genes associated with cellular stress response (*dksA*, *rpoS* and *uvrD*). Among the TCA cycle genes, *sdhB* and *sdhC* encode subunits of the electron transport chain enzyme succinate dehydrogenase complex and share an operon. Previous studies in *E. coli* and *Corynebacterium glutamicum* indicated that TCA cycle activity—particularly succinate dehydrogenase—supports lycopene production by increasing ATP and NADPH availability (Göttl et al., 2021; Sun et al., 2014). Among the cell stress response genes, prior work reported that disruption of *rpoS* expression decreased lycopene accumulation in *E. coli* independently of MEP pathway flux, due to increased reactive oxygen species during late-log and stationary phase that degrade the oxidation-sensitive lycopene molecule (Bongers et al., 2015). More generally, entry into stationary phase promotes metabolite availability for lycopene production (Farmer and Liao, 2001), and *rpoS* expression accordingly impacts lycopene yield (Becker-Hapak et al., 1997). Together, our results show that disruptions in the TCA cycle, the bacterial stress response, and core isoprenoid biosynthesis are deleterious to lycopene accumulation.

In contrast, repression of 18 genes across 5 cellular processes significantly increased lycopene yield (**Figure 4C,D**). Some of the largest increases in lycopene yield were associated with fatty acid biosynthesis genes *accA* and *accD* (which are among the green points in the top left corner of **Figure 4B**). However, repression of these genes also strongly reduced the final OD_600_. When analyzing the lycopene yield in units of milligrams per liter (rather than normalized by OD_600_), we observed that five of the six *acc*-gene targeting strains had either a decreased or no significant change in lycopene yield relative to the NT strain (**Table S5**). We also observed increased lycopene yield for the pyruvate fermentation gene *pflB* and several genes associated with the TCA cycle including *aceE*, *aceF*, *acnA*, and *sdhD*. Previous work combining flux balance analysis and experimental validation showed that repression of central carbon metabolism genes such as *aceE* can improve lycopene yield in glucose minimal media (Alper et al., 2005; Alper and Stephanopoulos, 2008). Our results further indicate a role for central carbon metabolism genes towards isoprenoid yield in LB medium, in the absence of glucose as a carbon source. We also observed lycopene yield increased for four genes in amino acid biosynthesis including branched chain amino acid synthesis enzymes *ilvD* and *leuC*, and chorismate biosynthesis enzymes *aroK* and *aroF*. To our knowledge, these four genes have not previously been implicated in metabolic engineering strategies for carotenoid production. Phospholipid biosynthesis genes *dgkA* and *pgsA* both increased lycopene yield when repressed. The gene *dgkA* has already been studied as a lycopene-modulating gene (Li et al., 2024; Wu et al., 2018). The gene *pgsA* encodes an enzyme that catalyzes the first committed step toward biosynthesis of acidic phospholipids. Finally, repression of several SOS response genes (*umuC*, *uvrB*, *uvrD*) and the *clpP* gene encoding a stringent response protease produced increases in lycopene yield, highlighting a possible link between the cell response to stress and carotenoid biosynthesis.

To summarize pathway-level patterns among the lycopene-modulating genes, we calculated the change in lycopene production of each strain relative to the NT control and averaged this effect across all sgRNAs targeting a given pathway (**Figure 4E**). The contributions of individual gene knockdowns to lycopene yield are shown in **Figure 4F**, to provide a sense of how the effect is distributed across a pathway. The pathway with the strongest increase in lycopene production is fatty acid biosynthesis driven by five of eight constituent genes, though these five strains were among the six poorest growing strains with respect to OD_600_ (**Figure 4B, Table S5**). The next strongest increase in lycopene production was for acetate metabolism with a single member gene *pta*. The gene *pta* is an acetyl transferase that catalyzes the reversible reaction of loading acetyl phosphate onto CoA making acetyl-CoA. Then the next three pathway groups are all close to this reaction: fermentation, glycolysis/gluconeogenesis, and glycolysis. Together, these results support the idea that reduced abundance of metabolic enzymes in pathways near pyruvate in central carbon metabolism reallocates pyruvate toward lycopene production. Four pathways had an average decrease in lycopene yield. Aspartate and gluconeogenesis pathway effects on lycopene are each driven by single genes *ascA* and *tpiA* respectively. While the lycopene yields of both trend toward decreasing, neither gene showed a significant decrease in lycopene production after false-discovery rate correction. The strongest effect was isoprenoid biosynthesis, which was expected due to the precursors of lycopene biosynthesis made by these enzymes. Overall, these results identify genes across diverse metabolic pathways that influence lycopene biosynthesis and highlight potential targets for engineering improved intracellular isoprenoid production.

## DISCUSSION AND CONCLUSIONS

Metabolic engineering often requires optimization of gene expression levels across both the heterologous pathway and host metabolism to reduce toxicity and increase flux. This requires a generalizable strategy to map expression-metabolic yield relationships throughout metabolism. To this end, we introduce a CRISPRi-based approach to identify *E. coli* genes as modulators of isoprenoid production. We developed a high yield lycopene-producing chassis strain as a starting point for our CRISPRi screen and quantified the impact of gene repression on lycopene production across 180 genes from 24 *E. coli* pathways. Importantly, our method integrates NGS to permit mapping of each CRISPRi sgRNA to lycopene phenotype. In total, we identified 19 strains where gene repression decreased lycopene production, and 18 strains where gene repression increased lycopene production.

Our screen was conducted in LB media, in contrast to several earlier studies that made use of M9 minimal media with glucose. However, like earlier work, we identified numerous genes in the TCA cycle and central carbon metabolism with a role in isoprenoid production, indicating that these genes play a key role across different culture conditions. Across the TCA cycle we found diverging roles for gene repression on lycopene yield: repression of *sdhC*, *sdhB*, *sucB*, and *acnB* decreased yield, while repression of *aceE*, *aceF*, *acnA*, and *sdhD* increased yield. This varied response to gene repression across the TCA cycle is likely a result of the complexities of metabolic flux networks, and/or may reflect polar effects of CRISPRi knockdowns on multi-gene operons. To explain, consider that the genes encoding the succinate dehydrogenase complex comprise the *sdhCDAB* operon. The *sdhCDAB* operon is located upstream to the sucAB operon and sometimes both *sucAB* and *sdhCDAB* are co-transcribed. Thus, using CRISPRi to target the first gene in the operon (*sdh*C) likely interferes with expression for all *sdh* and *suc* genes. In keeping with this, CRISPRi repression of *sdhC* had the strongest magnitude of reduced lycopene production across the targeted succinate dehydrogenase (*sdh*) and 2-oxoglutarate dehydrogenase complex (*suc*) genes. However, targeting the second gene in the operon (*sdhD*) yielded a significant *increase* in lycopene yield. The genes encoded by *sdhDAB* can form a separate transcriptional unit that is regulated by catabolite repressor protein (CRP). Additionally, *sdhB* is known to be preferentially expressed separately from *sdhC* depending on the state of the quinone pool regulating *arcB* sensor kinase (Alvarez et al., 2013; Sharma et al., 2013). Consequently, CRISPRi may interfere with expression (and metabolic outcomes) differently depending on sgRNA targeting within the operon and interactions with regulatory factors like *arcB* and CRP. These findings illustrate the potential for differential effects of gene knockdown across an operon, and encourage follow-up investigations studying *arcAB* sensor activity, changes to ubiquinone species in lycopene producing strains, and the repression efficiency of each guide targeting elements of the *sdh* operon.

We also identified several genes in amino acid metabolism (*ilvA*, *leuC*, *aroK*, and *aroF*) with roles in lycopene biosynthesis. In all cases, CRISPRi mediated repression increased lycopene production. The genes *ilvA* and *leuC* encode the two steps of isoleucine synthesis. The gene *aroF* catalyzes the first committed step of chorismate synthesis, while *aroK* encodes another chorismate biosynthesis enzyme; it also is the leading gene on an operon with many other metabolic enzymes. In LB media, there is a documented excess of amino acids, including branched chain amino acids, and isoleucine in particular. Through this lens, one possible explanation is that biosynthesis of branched chain amino acids is not necessary in LB culture conditions, and so repressing their synthetic machinery saves metabolites for other anabolic processes. To our knowledge, none of these four genes have previously been connected to lycopene production, though these findings may be specific to *E. coli* grown in LB media.

Together these results highlight the value in conducting a broad screen that samples genes across metabolism and cell regulation. By pairing inducible gene repression with medium-throughput phenotyping, we uncovered diverse modulators of isoprenoid yield across central carbon metabolism, amino acid biosynthesis, phospholipid production, membrane physiology, stress response pathways, and electron transport components, illustrating the multifaceted cellular constraints that shape isoprenoid yield. These findings highlight that beneficial tuning often arises not only from canonical precursor pathways but also from genes whose endogenous expression exceeds metabolic needs under LB culture conditions. While our current assay focuses on single-gene knockdowns, CRISPRi makes it possible to create combinatorial perturbations across multiple genes (Hawkins et al., 2020; Otto et al., 2024). This opens up the possibility of defining the pattern of genetic interactions (epistasis) that shape biosynthetic yield (Alper and Stephanopoulos, 2008). Moreover, because the library is arrayed into individual wells for characterization, this style of assay could be combined with other quantitative readouts of biochemical phenotype, including MALDI mass spectrometry or imaging. By combining such measurements with machine learning (Otto et al., 2024) one could estimate the effects of perturbing multiple genes in combination — in essence, constructing a high-dimensional expression-biosynthetic yield landscape. Together our work provides a foundation for future metabolic engineering efforts that integrate CRISPRi screens, environmental context, and modeling to optimize complex biosynthetic pathways.

## ACKNOWLEDGEMENTS

The authors thank members of the Reynolds and Saunders lab for early feedback on the project. We also thank Dr. Boyuan Wang for his input and suggestions on the manuscript. This work was supported in part by the Green Center for Systems Biology at UT Southwestern Medical Center, and the National Institute of General Medical Sciences of the National Institutes of Health under Award Number R01GM136842 to KAR.

## AUTHOR CONTRIBUTIONS

**Dhiraj Dokwal:** Conceptualization, Methodology, Formal analysis, Investigation, Writing – Original draft **Philip M. Brown**: Formal analysis, Investigation, Writing – Original draft, Visualization **Christine Ingle**: Investigation, Writing – Review & editing **Scott H. Saunders**: Methodology, Resources, Writing – Review & editing **Kimberly A. Reynolds**: Conceptualization, Formal analysis, Writing – Review & editing, Supervision, Funding acquisition

## Notes

### Competing Interest Statement

The authors have declared no competing interest.

