## Supplemental Information for "Systematic CRISPRi screening reveals genetic modulators of *E. coli* isoprenoid production"

**Contents:**

**I. Supplementary Figures**

**Fig. S1:** Genetics of the lycopene-producing *E. coli* chassis strain.

**Fig. S2**: The effect of CRISPRi induction time and cell growth on lycopene production.

**II. Supplementary Tables**

**Table S1:** Primers for cloning, sequencing, and next generation sequencing amplicon preparation.

**Table S2:** Focused sgRNA library targeting several key genes in lycopene biosynthesis. (Related to Figure 3).

**Table S3**: Large-scale sgRNA library targeting *E. coli* genes throughout metabolism. Annotations of target gene cell process, pathway, and essentiality (Related to Figure 4).

**Table S4**: Average and standard deviation of OD600 and Lycopene yield (mg/(L*OD600)) for each identified sgRNA (Related to Figure 4).

**Table S5**: Selected fatty acid biosynthesis genes from the large scale sgRNA library. Average and standard deviations of lycopene yield (mg/L), as well as p-values from two-tailed, unequal variance, t-test for each strain versus the NT control.

A

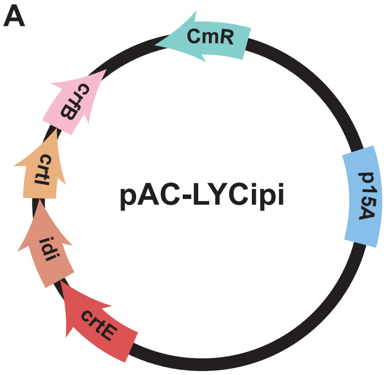


B

**Wild Type RBS**

TAAACAAT--AAGTATTAATAGGCCCCTGatgAGTTTT

TAAAAATTTCGATTACAGGAATTACGTTAatgAGTTTT

***dxs* 5000x RBS**

C

**Wild Type**

GCTTTTTATTCTGTCTCAACTCTGGATGTTTC--atgAAGCAA

GCTTTTTATTATACATCGAAACAAGGACGTATAAatgAAGCAA

***dxr* 6000x RBS**

**Fig. S1. Genetics of the lycopene-producing *E. coli* chassis strain.**

**A)** Schematic of the pAC-LYCipi plasmid encoding the heterologous lycopene synthesis genes**.**

**B)** Native (top row) and edited (bottom row) ribosome binding sequence (RBS) for *E. coli* gene *dxs***.** Edits around the RBS region are shown in red; green indicates the beginning of the *dxs* open reading frame.

**C)** Native (top row) and edited (bottom row) ribosome binding sequence (RBS) for *E. coli* gene *dxr***.** Edits around the RBS region are shown in red; green indicates the beginning of the *dxr* open reading frame.

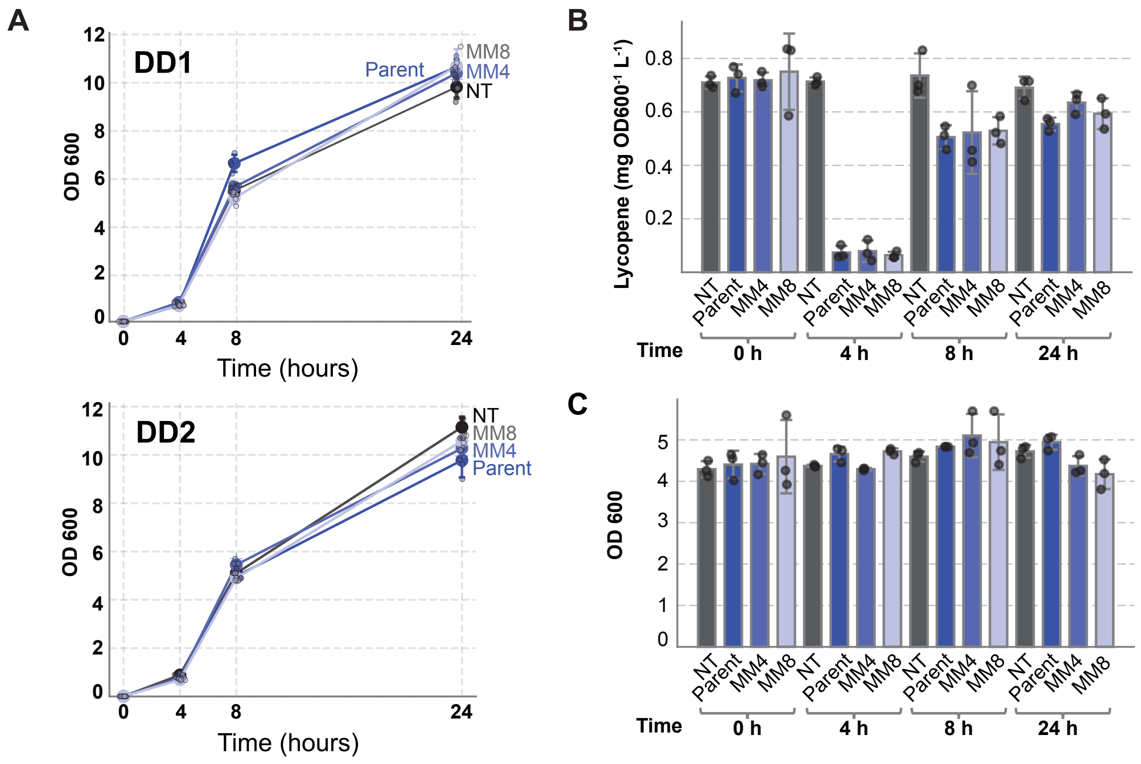


**Fig. S2.** The effect of CRISPRi induction time on lycopene production

**A)** OD600 measurements over time for strains DD1 and DD2 cultured in deep well plates. Each colored line indicates a distinct sgRNA plasmid. As in main text figure 2, the parent sgRNA targets *dxs* (dark blue line). MM4 (light blue) and MM8 (grey blue) are sgRNA variants derived from the *dxs*-targeting parent with 4 or 8 mismatches in the homology region respectively. NT (black line) indicates a non-targeting sgRNA. Each point indicates the mean over triplicate measurements; error bars represent standard deviation. CRISPRi was not induced during the collection of these growth curves.

**B)** Lycopene yield (mg OD600$^{-1}$ L$^{-1}$) of strain DD1 carrying the NT, Parent, MM4, or MM8 sgRNAs after CRISPRi induction at the given timepoints. Color coding is identical to panel (**A**). Points indicate the individual replicates (3 each), the bar heights reflect the mean across the triplicate measurements, and the error bars represent standard deviation.

**C)** Endpoint culture density (OD600) of strain DD1 carrying the NT, Parent, MM4, or MM8 sgRNAs after 48 hours of culture following CRISPRi induction at the given timepoints. Color coding is identical to panel (**A**). Points indicate individual replicates (3 per strain/sgRNA combination), bar height reflects the mean across the triplicate measurements, and the error bars represent standard deviation.

| **Name** | **Sequence** | **Reaction** |
|---|---|---|
| *dxs* RBS PCR screening forward primer | GCAGTTCGTCGCAGAGTTTC | Screening ORBIT RBS insertion |
| *dxs* RBS PCR screening reverse primer | TCCTGGATGTGGTGGGAGAT | Screening ORBIT RBS insertion |
| *dxr* RBS PCR screening forward primer | GTGACGCGAACGACAAAGTG | Screening ORBIT RBS insertion |
| *dxr* RBS PCR screening reverse primer | GTGCCTGGAATTCTCTCCCC | Screening ORBIT RBS insertion |
| *dxs* targeting oligo | GGACTACATCATCCAGCGTAATAAATAAggcttgtcgacgacggcggtctccgtcgtcaggatcat AAATTTCGATTACAGGAATTACGTTAATGAGTTTTGATATTGCCAAATACCCGA | ORBIT of RBS for *dxs* |
| *dxr* targeting oligo | GTCGAGCCCAGAATGGTGAGTTGCTTcatTTATACGTCCTTGTTTCGATGTATatgatcctgacgacggagaccgccgtcgtcgacaagccAATAAAAAGCAAAACGCCGCCAGCCGATC | ORBIT of RBS for *dxr* |
| GG_univ_F | GTACTGGGTCTCTAGGTTTGACAGCTAGCTCAGTCCTAG | Bsa-I overhang on oligo for golden gate cloning of sgRNAs |
| GG_univ_R | GTACTGGGTCTCTCCACCTTCAAAAAAAGCACCGACTCG | Bsa-I overhang on oligo for golden gate cloning of sgRNAs |
| Truseq_F1 | TGACTGGAGTTCAGACGTGTGCTCTTCCGATCTNNNNAATAGGCGCTTCACCTCGAGAGGTTTGACAG | Amplicon generation: Truseq |

| | | |
|---|---|---|
| | | adapter addition and plate one barcode |
| Truseq_F2 | TGACTGGAGTTCAGACGTGTGCTCTTCCGATCTNNNNAATGAGCCCTTCACCTCGAGAGGTTTGACAG | Amplicon generation: Truseq adapter addition and plate two barcode |
| Truseq_F3 | TGACTGGAGTTCAGACGTGTGCTCTTCCGATCTNNNNAATCTCCGCTTCACCTCGAGAGGTTTGACAG | Amplicon generation: Truseq adapter addition and plate three barcode |
| Truseq_F4 | TGACTGGAGTTCAGACGTGTGCTCTTCCGATCTNNNNAAGAAGGCCTTCACCTCGAGAGGTTTGACAG | Amplicon generation: Truseq adapter addition and plate four barcode |
| Truseq_F | TGACTGGAGTTCAGACGTGTGCTCTTCCGATCTNNNNCTTCACCTCGAGAGGTTTGACAG | Truseq adapter addition |
| TruSeq_R | CACTCTTTCCCTACACGACGCTCTTCCGATCTNNNNNCCATTATTAGTACAGCGAGGCAAC | Truseq adapter addition |
| i5 Primer | AATGATACGGCGACCACCGAGATCTACACXXXXXXXXACACTCTTTCCCTACACGACGCTCTTCCGATCT | Illumina Index addition - 8 for rows |
| i7 Primer | CAAGCAGAAGACGGCATACGAGATXXXXXXXXGTGACTGGAGTTCAGACGTGTGCTCTTCCGATC | Illumina Index addition - 12 for columns |

**Table S1:** Primers for cloning, sequencing, and next generation sequencing amplicon preparation. Barcode regions are indicated in underline. In this table “N” indicates randomized nucleotides (mix of A/C/T/G), and “X” indicates the barcode region of the Illumina index adaptors. We used the standard TruSeq A501-508 and A701-712 barcodes from Illumina.

| sgRNA Name | Homology Sequence | Average Lycopene (mg/(OD600*L)) | Standard Deviation Lycopene | Number of Replicates |
|---|---|---|---|---|
| dxr_1_14_C | CGAGCCGGTCGAGCCCAGAA | | | 0 |
| dxr_1_14_MM2 | GCAGCCGGTCGAGCCCAGAA | | | 0 |
| dxr_1_14_MM4 | GCTCCCGGTCGAGCCCAGAA | 0.094 | | 1 |
| dxr_1_14_MM6 | GCTCGGGGTCGAGCCCAGAA | | | 0 |
| dxr_1_14_MM8 | GCTCGGCCTCGAGCCCAGAA | | | 0 |
| dxr_1_14_MM10 | GCTCGGCCAGGAGCCCAGAA | | | 0 |
| dxr_1_14_MM12 | GCTCGGCCAGCTGCCCAGAA | 0.210 | | 1 |
| dxr_1_14_MM14 | GCTCGGCCAGCTCGCCAGAA | | | 0 |
| dxr_3_222_C | GCCCACTTAAGACTTCGGTG | 0.011 | 0.01 | 2 |
| dxr_3_222_MM2 | CGCCACTTAAGACTTCGGTG | | | 0 |
| dxr_3_222_MM4 | CGGGACTTAAGACTTCGGTG | | | 0 |
| dxr_3_222_MM6 | CGGGTGTTAAGACTTCGGTG | | | 0 |
| dxr_3_222_MM8 | CGGGTGAAAAGACTTCGGTG | | | 0 |
| dxr_3_222_MM10 | CGGGTGAATTGACTTCGGTG | | | 0 |
| dxr_3_222_MM12 | CGGGTGAATTCTCTTCGGTG | | | 0 |
| dxr_3_222_MM14 | CGGGTGAATTCTGATCGGTG | 0.248 | 0.016 | 2 |
| ispA_1_10_C | ACGCAGGCTTCGAGTTGCTG | | | 0 |
| ispA_1_10_MM2 | TGGCAGGCTTCGAGTTGCTG | | | 0 |
| ispA_1_10_MM4 | TGCGAGGCTTCGAGTTGCTG | | | 0 |
| ispA_1_10_MM6 | TGCGTCGCTTCGAGTTGCTG | | | 0 |
| ispA_1_10_MM8 | TGCGTCCGTTCGAGTTGCTG | 0.225 | | 1 |
| ispA_1_10_MM10 | TGCGTCCGAACGAGTTGCTG | 0.252 | | 1 |
| ispA_1_10_MM12 | TGCGTCCGAAGCAGTTGCTG | 0.237 | | 1 |
| ispA_1_10_MM14 | TGCGTCCGAAGCTCTTGCTG | | | 0 |
| ispA_2_68_C | AGTGTTCTGAAAGGGCAGTG | | | 0 |
| ispA_2_68_MM2 | TCTGTTCTGAAAGGGCAGTG | 0.104 | 0.082 | 2 |
| ispA_2_68_MM4 | TCACTTCTGAAAGGGCAGTG | 0.089 | | 1 |
| ispA_2_68_MM6 | TCACAACTGAAAGGGCAGTG | 0.074 | | 1 |
| ispA_2_68_MM8 | TCACAAGAGAAAGGGCAGTG | | | 0 |
| ispA_2_68_MM10 | TCACAAGACTAAGGGCAGTG | | | 0 |
| ispA_2_68_MM12 | TCACAAGACTTTGGGCAGTG | | | 0 |
| ispA_2_68_MM14 | TCACAAGACTTTCCGCAGTG | 0.231 | | 1 |
| ldhA_1_45_C | AGGACTCGTTCACCTGTTGC | 0.281 | 0.07 | 2 |
| ldhA_1_45_MM2 | TCGACTCGTTCACCTGTTGC | | | 0 |
| ldhA_1_45_MM4 | TCCTCTCGTTCACCTGTTGC | | | 0 |

| | | | | |
|---|---|---|---|---|
| ldhA_1_45_MM6 | TCCTGACGTTCACCTGTTGC | 0.249 | | 1 |
| ldhA_1_45_MM8 | TCCTGAGCTTCACCTGTTGC | 0.221 | 0.093 | 4 |
| ldhA_1_45_MM10 | TCCTGAGCAACACCTGTTGC | 0.246 | 0.098 | 3 |
| ldhA_1_45_MM12 | TCCTGAGCAAGTCCTGTTGC | 0.242 | 0.062 | 2 |
| ldhA_1_45_MM14 | TCCTGAGCAAGTGGTGTTGC | | | 0 |
| ldhA_2_125_C | ACATACCGCTTCGCAGCCAT | | | 0 |
| ldhA_2_125_MM2 | TGATACCGCTTCGCAGCCAT | 0.261 | | 1 |
| ldhA_2_125_MM4 | TGTAACCGCTTCGCAGCCAT | | | 0 |
| ldhA_2_125_MM6 | TGTATGCGCTTCGCAGCCAT | | | 0 |
| ldhA_2_125_MM8 | TGTATGGCCTTCGCAGCCAT | 0.234 | | 1 |
| ldhA_2_125_MM10 | TGTATGGCGATCGCAGCCAT | | | 0 |
| ldhA_2_125_MM12 | TGTATGGCGAAGGCAGCCAT | 0.303 | 0.073 | 2 |
| ldhA_2_125_MM14 | TGTATGGCGAAGCGAGCCAT | 0.247 | | 1 |
| dxs_1_17_C | CAGTGCCAGGGTCGGGTATT | 0.005 | | 1 |
| dxs_1_17_MM2 | GTGTGCCAGGGTCGGGTATT | 0.018 | | 1 |
| dxs_1_17_MM4 | GTCAGCCAGGGTCGGGTATT | | | 0 |
| dxs_1_17_MM6 | GTCACGCAGGGTCGGGTATT | | | 0 |
| dxs_1_17_MM8 | GTCACGGTGGGTCGGGTATT | 0.011 | | 1 |
| dxs_1_17_MM10 | GTCACGGTCCGTCGGGTATT | 0.062 | | 1 |
| dxs_1_17_MM12 | GTCACGGTCCCACGGGTATT | | | 0 |
| dxs_1_17_MM14 | GTCACGGTCCCAGCGGTATT | | | 0 |
| dxs_3_216_C | GCCCCACATCCCAAATCAAT | 0.024 | 0.001 | 2 |
| dxs_3_216_MM2 | CGCCCACATCCCAAATCAAT | 0.046 | | 1 |
| dxs_3_216_MM4 | CGGGCACATCCCAAATCAAT | | | 0 |
| dxs_3_216_MM6 | CGGGGTCATCCCAAATCAAT | 0.024 | 0.01 | 2 |
| dxs_3_216_MM8 | CGGGGTGTTCCCAAATCAAT | 0.018 | | 1 |
| dxs_3_216_MM10 | CGGGGTGTAGCCAAATCAAT | 0.170 | | 1 |
| dxs_3_216_MM12 | CGGGGTGTAGGGAAATCAAT | | | 0 |
| dxs_3_216_MM14 | CGGGGTGTAGGGTTATCAAT | 0.321 | | 1 |
| gdhA_1_42 | TTTGATTCGGGTCGCGCTTT | | | 0 |
| dxr_2_66_C | CGCGGAAGTGTTCGGGATTA | | | 0 |
| dxr_2_66_MM2 | GCCGGAAGTGTTCGGGATTA | | | 0 |
| dxr_2_66_MM4 | GCGCGAAGTGTTCGGGATTA | | | 0 |
| dxr_2_66_MM6 | GCGCCTAGTGTTCGGGATTA | | | 0 |
| dxr_2_66_MM8 | GCGCCTTCTGTTCGGGATTA | 0.100 | 0.048 | 2 |
| dxr_2_66_MM10 | GCGCCTTCACTTCGGGATTA | 0.160 | 0.119 | 2 |
| dxr_2_66_MM12 | GCGCCTTCACAACGGGATTA | 0.086 | | 1 |
| dxr_2_66_MM14 | GCGCCTTCACAAGCGGATTA | 0.219 | 0.05 | 3 |
| ispA_3_205_C | ATACACTCAACGGCGGCAGC | | | 0 |
| ispA_3_205_MM2 | TAACACTCAACGGCGGCAGC | 0.110 | | 1 |
| ispA_3_205_MM4 | TATGACTCAACGGCGGCAGC | 0.138 | | 1 |

| | | | | |
|---|---|---|---|---|
| ispA_3_205_MM6 | TATGTGTCAACGGCGGCAGC | | | 0 |
| ispA_3_205_MM8 | TATGTGAGAACGGCGGCAGC | | | 0 |
| ispA_3_205_MM10 | TATGTGAGTTCGGCGGCAGC | | | 0 |
| ispA_3_205_MM12 | TATGTGAGTTGCGCGGCAGC | 0.265 | 0.013 | 2 |
| ispA_3_205_MM14 | TATGTGAGTTGCCGGGCAGC | | | 0 |
| ldhA_3_218_C | ATTGAAACCGGCACAGCGCA | 0.292 | | 1 |
| ldhA_3_218_MM2 | TATGAAACCGGCACAGCGCA | | | 0 |
| ldhA_3_218_MM4 | TAACAAACCGGCACAGCGCA | | | 0 |
| ldhA_3_218_MM6 | TAACTTACCGGCACAGCGCA | | | 0 |
| ldhA_3_218_MM8 | TAACTTTGCGGCACAGCGCA | | | 0 |
| ldhA_3_218_MM10 | TAACTTTGGCGCACAGCGCA | | | 0 |
| ldhA_3_218_MM12 | TAACTTTGGCCGACAGCGCA | 0.235 | | 1 |
| ldhA_3_218_MM14 | TAACTTTGGCCGTGAGCGCA | 0.228 | | 1 |
| dxs_2_47_C | CAACAGTCGTAACTCCTGGG | | | 0 |
| dxs_2_47_MM2 | GTACAGTCGTAACTCCTGGG | 0.010 | | 1 |
| dxs_2_47_MM4 | GTTGAGTCGTAACTCCTGGG | | | 0 |
| dxs_2_47_MM6 | GTTGTCTCGTAACTCCTGGG | | | 0 |
| dxs_2_47_MM8 | GTTGTCAGGTAACTCCTGGG | 0.008 | | 1 |
| dxs_2_47_MM10 | GTTGTCAGCAAACTCCTGGG | | | 0 |
| dxs_2_47_MM12 | GTTGTCAGCATTCTCCTGGG | | | 0 |
| dxs_2_47_MM14 | GTTGTCAGCATTGACCTGGG | 0.302 | 0.15 | 2 |
| NT (negC_rand_42) | AGTAATTGTATAGGCGACCT | 0.198 | | |
| Parent sgRNA | | 0.015 | | |
| HW2e | | 0.203 | | |
| HW2f | | 0.168 | | |

**Table S2:** Focused sgRNA library targeting several key genes in lycopene biosynthesis. (Related to Figure 3).

| Guide | Pathway | Category | Homology Sequence | Essential | Recovered |
|---|---|---|---|---|---|
| negC_rand_42 | N.D. | N.D. | AGTAATTGTA TAGGCGACCT | 0 | 1 |
| acnB_1_15_C | TCA cycle | Central Carbon Metabolism | CACGCTCAGC TACGTGCTTA | 0 | 1 |
| acnB_1_15_MM 7 | TCA cycle | Central Carbon Metabolism | GTGCGAGAG CTACGTGCTT A | 0 | 1 |
| dxr_1_14_C | Isoprenoid Biosynthesis | Ubiquinone Biosynthesis | CGAGCCGGT CGAGCCCAGA A | 1 | 0 |
| dxr_1_14_MM7 | Isoprenoid Biosynthesis | Ubiquinone Biosynthesis | GCTCGGCGTC GAGCCCAGAA | 1 | 1 |
| fadB_1_20_C | β-oxidation | Fatty Acid Metabolism | TTCCAGCCAG TCAAGGTACA | 0 | 0 |
| fadB_1_20_MM 7 | β-oxidation | Fatty Acid Metabolism | AAGGTCGCAG TCAAGGTACA | 0 | 0 |
| dnaA_1_32_C | Stringent response | Stress Responses | TGGTAACTCA TCCTGCAATC | 1 | 0 |
| dnaA_1_32_MM 7 | Stringent response | Stress Responses | ACCATTGTCA TCCTGCAATC | 1 | 0 |
| dksA_1_32_C | Stringent response | Stress Responses | AGCGATGGC GAGAATACTC A | 0 | 1 |

| | | | | | |
|---|---|---|---|---|---|
| dksA_1_32_MM7 | Stringent response | Stress Responses | TCGCTACGCGAGAATACTCA | 0 | 1 |
| ispD_1_8_C | Isoprenoid Biosynthesis | Ubiquinone Biosynthesis | GGCGCAAACATCCAAATGAG | 1 | 1 |
| ispD_1_8_MM7 | Isoprenoid Biosynthesis | Ubiquinone Biosynthesis | CCGCGTTACATCCAAATGAG | 1 | 1 |
| fadH_1_9_C | β-oxidation | Fatty Acid Metabolism | CCAGCGGGGCGAACAGCGAC | 0 | 0 |
| fadH_1_9_MM7 | β-oxidation | Fatty Acid Metabolism | GGTCGCCGGCGAACAGCGAC | 0 | 1 |
| ispA_1_10_C | Isoprenoid Biosynthesis | Ubiquinone Biosynthesis | ACGCAGGCTTCGAGTTGCTG | 1 | 1 |
| ispA_1_10_MM7 | Isoprenoid Biosynthesis | Ubiquinone Biosynthesis | TGCGTCCCTTCGAGTTGCTG | 1 | 1 |
| sdhD_1_14_C | TCA cycle | Central Carbon Metabolism | GCCATTGCGTCCTAATGCGG | 0 | 0 |
| sdhD_1_14_MM7 | TCA cycle | Central Carbon Metabolism | CGGTAACCGTCCTAATGCGG | 0 | 1 |
| folP_1_14_C | Folate Biosynthesis | Folate Biosynthesis | AAGGTCCAGTGAAGTACCCT | 1 | 1 |
| folP_1_14_MM7 | Folate Biosynthesis | Folate Biosynthesis | TTCCAGGAGTGAAGTACCCT | 1 | 0 |

| | | | | | |
|---|---|---|---|---|---|
| aroD_1_83_C | Chorismate Biosynthesis | Amino Acid Biosynthesis | GAGAGCTTCG GATTTCACGC | 0 | 1 |
| aroD_1_83_MM 7 | Chorismate Biosynthesis | Amino Acid Biosynthesis | CTCTCGATCG GATTTCACGC | 0 | 0 |
| ubiG_1_27_C | ubiquinone (coenzyme Q) biosynthesis | Ubiquinone Biosynthesis | TCTCTTCGTG GTCTACGTTA | 1 | 0 |
| ubiG_1_27_MM 7 | ubiquinone (coenzyme Q) biosynthesis | Ubiquinone Biosynthesis | AGAGAAGGTG GTCTACGTTA | 1 | 0 |
| trpE_1_16_C | Stringent response | Stress Responses | CAGGTTAGCA GTTCGAGAGT | 0 | 0 |
| trpE_1_16_MM7 | Stringent response | Stress Responses | GTCCAATGCA GTTCGAGAGT | 0 | 0 |
| ilvA_1_16_C | Branched Chain Amino Acid Metabolism | Amino Acid Biosynthesis | CCTTCCGGAG CACCGGACAG | 0 | 1 |
| ilvA_1_16_MM7 | Branched Chain Amino Acid Metabolism | Amino Acid Biosynthesis | GGAAGGCGA GCACCGGACA G | 0 | 0 |
| sucA_1_23_C | TCA cycle | Central Carbon Metabolism | GAGGTAAGAA GAGTCCAACC | 1 | 0 |
| sucA_1_23_MM 7 | TCA cycle | Central Carbon Metabolism | CTCCATTGAA GAGTCCAACC | 1 | 0 |
| maeA_1_39_C | Malate metabolism | Central Carbon Metabolism | GCAGTACAGG GCCAGCGTAA | 0 | 0 |

| | | | | | |
|---|---|---|---|---|---|
| maeA_1_39_MM7 | Malate metabolism | Central Carbon Metabolism | CGTCATGAGGGCCAGCGTAA | 0 | 0 |
| yajO_1_26_C | Isoprenoid Biosynthesis | Ubiquinone Biosynthesis | AAGTCGGGAAACGCGAAGGT | 0 | 1 |
| yajO_1_26_MM7 | Isoprenoid Biosynthesis | Ubiquinone Biosynthesis | TTCAGCCGAAACGCGAAGGT | 0 | 1 |
| aroA_1_8_C | Chorismate Biosynthesis | Amino Acid Biosynthesis | AGCGATGGGTTGTAACGTCA | 0 | 1 |
| aroA_1_8_MM7 | Chorismate Biosynthesis | Amino Acid Biosynthesis | TCGCTACGGTTGTAACGTCA | 0 | 0 |
| polA_1_76_C | SOS response | Stress Responses | CCTGCGCTGTTAGTCAGCGG | 0 | 1 |
| polA_1_76_MM7 | SOS response | Stress Responses | GGACGCGTGTTAGTCAGCGG | 0 | 1 |
| pck_1_23_C | Gluconeogenesis | Central Carbon Metabolism | ATAAGCCTCGAGTTCTTGCG | 0 | 0 |
| pck_1_23_MM7 | Gluconeogenesis | Central Carbon Metabolism | TATTCGGTCGAGTTCTTGCG | 0 | 1 |
| aceF_1_22_C | TCA cycle | Central Carbon Metabolism | ACTTCATCAGCCCCGATGTC | 1 | 0 |
| aceF_1_22_MM7 | TCA cycle | Central Carbon Metabolism | TGAAGTACAGCCCCGATGTC | 1 | 1 |

| | | | | | |
|---|---|---|---|---|---|
| fbaB_1_70_C | Glycolysis | Central Carbon Metabolism | GGGAGATAAAGCTGGTCAGA | 0 | 1 |
| fbaB_1_70_MM7 | Glycolysis | Central Carbon Metabolism | CCCTCTAAAAGCTGGTCAGA | 0 | 0 |
| gpsA_1_38_C | Phospholipid/cardiolipin biosynthesis | Fatty Acid Metabolism | AAGAGCGGTGCCGTACGAGC | 1 | 1 |
| gpsA_1_38_MM7 | Phospholipid/cardiolipin biosynthesis | Fatty Acid Metabolism | TTCTCGCGTGCCGTACGAGC | 1 | 0 |
| fumB_1_14_C | TCA cycle | Central Carbon Metabolism | GAAAGGTGCCTGGTAGATAA | 0 | 0 |
| fumB_1_14_MM7 | TCA cycle | Central Carbon Metabolism | CTTTCCAGCCTGGTAGATAA | 0 | 0 |
| icd_1_22_C | TCA cycle | Central Carbon Metabolism | GTGATCTTCTTGCCTTGTGC | 0 | 0 |
| icd_1_22_MM7 | TCA cycle | Central Carbon Metabolism | CACTAGATCTTGCCTTGTGC | 0 | 1 |
| adhE_1_56_C | Fermentation | Central Carbon Metabolism | GAAACTGGCATATTCACGCT | 0 | 1 |
| adhE_1_56_MM7 | Fermentation | Central Carbon Metabolism | CTTTGACGCATATTCACGCT | 0 | 1 |
| pabB_1_68_C | Folate Biosynthesis | Folate Biosynthesis | CCACGGCAGGTGGCTTAAGC | 0 | 1 |

| | | | | | |
|---|---|---|---|---|---|
| pabB_1_68_MM7 | Folate Biosynthesis | Folate Biosynthesis | GGTGCCGAGGTGGCTTAAGC | 0 | 0 |
| mutH_1_5_C | SOS response | Stress Responses | AGAGAGCAGTGGGCGAGGTT | 0 | 0 |
| mutH_1_5_MM7 | SOS response | Stress Responses | TCTCTCGAGTGGGCGAGGTT | 0 | 0 |
| fadL_1_15_C | β-oxidation | Fatty Acid Metabolism | CGAGAGCAGACTTTGTAAAC | 0 | 0 |
| fadL_1_15_MM7 | β-oxidation | Fatty Acid Metabolism | GCTCTCGAGACTTTGTAAAC | 0 | 0 |
| rpoD_1_12_C | Stringent response | Stress Responses | GAAGTTTCAGCTGTGACTGC | 1 | 0 |
| rpoD_1_12_MM7 | Stringent response | Stress Responses | CTTCAAACAGCTGTGACTGC | 1 | 0 |
| pykF_1_179_C | Glycolysis | Central Carbon Metabolism | GGTATCAAGCAGGATAGCGG | 0 | 0 |
| pykF_1_179_MM7 | Glycolysis | Central Carbon Metabolism | CCATAGTAGCAGGATAGCGG | 0 | 0 |
| sdhA_1_44_C | TCA cycle | Central Carbon Metabolism | CGCGCGCATACCTGCGCCAC | 0 | 1 |
| sdhA_1_44_MM7 | TCA cycle | Central Carbon Metabolism | GCGCGCGATACCTGCGCCAC | 0 | 0 |

| | | | | | |
|---|---|---|---|---|---|
| aroK_1_128_C | Chorismate Biosynthesis | Amino Acid Biosynthesis | AACCCAGCCCACATCAGCTC | 0 | 1 |
| aroK_1_128_MM7 | Chorismate Biosynthesis | Amino Acid Biosynthesis | TTGGGTCCCCACATCAGCTC | 0 | 1 |
| rpoS_1_86_C | Stringent response | Stress Responses | ACTGGGTTCCTGTTCTACTA | 0 | 0 |
| rpoS_1_86_MM7 | Stringent response | Stress Responses | TGACCCATCCTGTTCTACTA | 0 | 1 |
| menA_1_21_C | Vitamin K biosynthesis | Vitamin Metabolism | TTTCCAGCCACGCCTGAGTT | 0 | 0 |
| menA_1_21_MM7 | Vitamin K biosynthesis | Vitamin Metabolism | AAAGGTCCCACGCCTGAGTT | 0 | 0 |
| ilvH_1_71_C | Branched Chain Amino Acid Metabolism | Amino Acid Biosynthesis | TTCAATGTTGTAGCCACGCT | 0 | 0 |
| ilvH_1_71_MM7 | Branched Chain Amino Acid Metabolism | Amino Acid Biosynthesis | AAGTTACTTGTAGCCACGCT | 0 | 0 |
| gpmA_1_60_C | Glycolysis | Central Carbon Metabolism | CGTCGTACCAACCGGTGAAA | 0 | 1 |
| gpmA_1_60_MM7 | Glycolysis | Central Carbon Metabolism | GCAGCATCCAACCGGTGAAA | 0 | 0 |
| gpp_1_8_C | Stringent response | Stress Responses | GGCTGCATACAGCGACGAGG | 0 | 1 |

| | | | | | |
|---|---|---|---|---|---|
| gpp_1_8_MM7 | Stringent response | Stress Responses | CCGACGTTACAGCGACGAGG | 0 | 1 |
| accB_1_99_C | Fatty Acid Biosynthesis | Fatty Acid Metabolism | AACTTGCGGCAGGAGCTGCA | 1 | 1 |
| accB_1_99_MM7 | Fatty Acid Biosynthesis | Fatty Acid Metabolism | TTGAACGGGCAGGAGCTGCA | 1 | 1 |
| talA_1_32_C | Pentose phosphate pathway | Pentose Phosphate Pathway | GCCGCTGTCTGCCACGACAG | 0 | 0 |
| talA_1_32_MM7 | Pentose phosphate pathway | Pentose Phosphate Pathway | CGGCGACTCTGCCACGACAG | 0 | 1 |
| ytjC_1_15_C | Glycolysis/gluconeogensis | Central Carbon Metabolism | GCGTTTCACCGTGGCGGACT | 0 | 1 |
| ytjC_1_15_MM7 | Glycolysis/gluconeogensis | Central Carbon Metabolism | CGCAAAGACCGTGGCGGACT | 0 | 1 |
| aceA_1_52_C | TCA cycle | Central Carbon Metabolism | CGAGTAATGCCTTCCCAACG | 0 | 1 |
| aceA_1_52_MM7 | TCA cycle | Central Carbon Metabolism | GCTCATTTGCCTTCCCAACG | 0 | 0 |
| sulA_1_89_C | SOS response | Stress Responses | GACAACTTCACTGATAAGCC | 0 | 1 |
| sulA_1_89_MM7 | SOS response | Stress Responses | CTGTTGATCACTGATAAGCC | 0 | 0 |

| | | | | | |
|---|---|---|---|---|---|
| clsA_1_35_C | Phospholipid/cardiolipin biosynthesis | Fatty Acid Metabolism | GAGCAACCAGTATCCCAGAA | 0 | 0 |
| clsA_1_35_MM7 | Phospholipid/cardiolipin biosynthesis | Fatty Acid Metabolism | CTCGTTGCAGTATCCCAGAA | 0 | 1 |
| ispU_1_40_C | Isoprenoid Biosynthesis | Ubiquinone Biosynthesis | ACATGACGGCAGCCATGCGC | 1 | 1 |
| ispU_1_40_MM7 | Isoprenoid Biosynthesis | Ubiquinone Biosynthesis | TGTACTGGGCAGCCATGCGC | 1 | 0 |
| uvrA_1_54_C | SOS response | Stress Responses | GCTTGTCGCGGGGGATAACG | 0 | 0 |
| uvrA_1_54_MM7 | SOS response | Stress Responses | CGAACAGGCGGGGGATAACG | 0 | 1 |
| lpd_1_45_C | TCA cycle | Central Carbon Metabolism | AGGCAGCGGAGTAACCTGCG | 1 | 1 |
| lpd_1_45_MM7 | TCA cycle | Central Carbon Metabolism | TCCGTCGGGAGTAACCTGCG | 1 | 1 |
| pgsA_1_15_C | Phospholipid/cardiolipin biosynthesis | Fatty Acid Metabolism | GGAACAGTGTAAGCAACGTA | 1 | 1 |
| pgsA_1_15_MM7 | Phospholipid/cardiolipin biosynthesis | Fatty Acid Metabolism | CCTTGTCTGTAAGCAACGTA | 1 | 1 |
| rpe_1_20_C | Pentose phosphate pathway | Pentose Phosphate Pathway | ATCAGCCGACAGAATTGAGG | 1 | 0 |

| | | | | | |
|---|---|---|---|---|---|
| rpe_1_20_MM7 | Pentose phosphate pathway | Pentose Phosphate Pathway | TAGTCGGGAC AGAATTGAGG | 1 | 1 |
| ldhA_1_45_C | Fermentation | Central Carbon Metabolism | AGGACTCGTT CACCTGTTGC | 0 | 1 |
| ldhA_1_45_MM7 | Fermentation | Central Carbon Metabolism | TCCTGAGGTT CACCTGTTGC | 0 | 0 |
| entA_1_32_C | Iron acquisition | Iron Acquisition | GCCGATACCT TTACCTGCGC | 0 | 1 |
| entA_1_32_MM 7 | Iron acquisition | Iron Acquisition | CGGCTATCCT TTACCTGCGC | 0 | 1 |
| ispG_1_9_C | Isoprenoid Biosynthesis | Ubiquinone Biosynthesis | TTCTACGTTG AATTGGAGCC | 1 | 0 |
| ispG_1_9_MM7 | Isoprenoid Biosynthesis | Ubiquinone Biosynthesis | AAGATGCTTG AATTGGAGCC | 1 | 1 |
| pgi_1_16_C | Glycolysis/glucone ogensis | Central Carbon Metabolism | TGCCAGGCAG CGGTCTGCGT | 0 | 0 |
| pgi_1_16_MM7 | Glycolysis/glucone ogensis | Central Carbon Metabolism | ACGGTCCCAG CGGTCTGCGT | 0 | 1 |
| tyrB_1_20_C | Branched Chain Amino Acid Metabolism | Amino Acid Biosynthesis | AAGAATCGGG TCGCCAGCGT | 0 | 0 |
| tyrB_1_20_MM7 | Branched Chain Amino Acid Metabolism | Amino Acid Biosynthesis | TTCTTAGGGG TCGCCAGCGT | 0 | 0 |

| | | | | | |
|---|---|---|---|---|---|
| aroC_1_38_C | Chorismate Biosynthesis | Amino Acid Biosynthesis | CCCGTGCGATTCGCCGAAGG | 0 | 0 |
| aroC_1_38_MM7 | Chorismate Biosynthesis | Amino Acid Biosynthesis | GGGCACGGATTCGCCGAAGG | 0 | 1 |
| ilvM_1_87_C | Branched Chain Amino Acid Metabolism | Amino Acid Biosynthesis | CCATATTCATTGAGCAGACG | 0 | 0 |
| ilvM_1_87_MM7 | Branched Chain Amino Acid Metabolism | Amino Acid Biosynthesis | GGTATAACATTGAGCAGACG | 0 | 0 |
| ilvN_1_41_C | Branched Chain Amino Acid Metabolism | Amino Acid Biosynthesis | TACGCCCGGATGGTTGCGAA | 0 | 0 |
| ilvN_1_41_MM7 | Branched Chain Amino Acid Metabolism | Amino Acid Biosynthesis | ATGCGGGGGATGGTTGCGAA | 0 | 0 |
| rpiB_1_104_C | Pentose phosphate pathway | Pentose Phosphate Pathway | ATCAGTACGCTCTGACGACC | 0 | 1 |
| rpiB_1_104_MM7 | Pentose phosphate pathway | Pentose Phosphate Pathway | TAGTCATCGCTCTGACGACC | 0 | 1 |
| pssA_1_39_C | Phospholipid/cardiolipin biosynthesis | Fatty Acid Metabolism | AAATCTTGGGTAGTTGGGCA | 1 | 1 |
| pssA_1_39_MM7 | Phospholipid/cardiolipin biosynthesis | Fatty Acid Metabolism | TTTAGAAGGGTAGTTGGGCA | 1 | 1 |
| ptsG_1_55_C | Glycolysis | Central Carbon Metabolism | GCGATAGGCAGTACGGATAC | 0 | 0 |

| | | | | | |
|---|---|---|---|---|---|
| ptsG_1_55_MM7 | Glycolysis | Central Carbon Metabolism | CGCTATCGCAGTACGGATAC | 0 | 0 |
| ilvD_1_4_C | Branched Chain Amino Acid Metabolism | Amino Acid Biosynthesis | GTGGTGGCGGAACGGTACTT | 0 | 1 |
| ilvD_1_4_MM7 | Branched Chain Amino Acid Metabolism | Amino Acid Biosynthesis | CACCACCCGGAACGGTACTT | 0 | 0 |
| sucC_1_38_C | TCA cycle | Central Carbon Metabolism | CGGTGCTGGTAAGCCATAGC | 0 | 1 |
| sucC_1_38_MM7 | TCA cycle | Central Carbon Metabolism | GCCACGAGGTAAGCCATAGC | 0 | 0 |
| accC_1_23_C | Fatty Acid Biosynthesis | Fatty Acid Metabolism | CAATGCAATCTCGCCGCGGT | 1 | 0 |
| accC_1_23_MM7 | Fatty Acid Biosynthesis | Fatty Acid Metabolism | GTTACGTATCTCGCCGCGGT | 1 | 0 |
| recR_1_8_C | SOS response | Stress Responses | AAGCTGTGTTAACAGCGGGC | 0 | 0 |
| recR_1_8_MM7 | SOS response | Stress Responses | TTCGACAGTTAACAGCGGGC | 0 | 0 |
| relA_1_80_C | Stringent response | Stress Responses | GCACTCACACGACTTCTGGC | 0 | 1 |
| relA_1_80_MM7 | Stringent response | Stress Responses | CGTGAGTCACGACTTCTGGC | 0 | 0 |

| | | | | | |
|---|---|---|---|---|---|
| fumC_1_48_C | TCA cycle | Central Carbon Metabolism | CGCCCCACAGCTTATCTGCC | 0 | 1 |
| fumC_1_48_MM7 | TCA cycle | Central Carbon Metabolism | GCGGGGTCAGCTTATCTGCC | 0 | 0 |
| pgk_1_20_C | Glycolysis | Central Carbon Metabolism | TTTCCCAGCAAGATCCAGAT | 1 | 0 |
| pgk_1_20_MM7 | Glycolysis | Central Carbon Metabolism | AAAGGGTGCAAGATCCAGAT | 1 | 1 |
| ilvE_1_120_C | Branched Chain Amino Acid Metabolism | Amino Acid Biosynthesis | CTTTGTGCGAGTCGTAGCAA | 0 | 1 |
| ilvE_1_120_MM7 | Branched Chain Amino Acid Metabolism | Amino Acid Biosynthesis | GAAACACCGAGTCGTAGCAA | 0 | 0 |
| clsC_1_4_C | Phospholipid/cardiolipin biosynthesis | Fatty Acid Metabolism | AGCACCGCGCTCGCCAGCCG | 0 | 1 |
| clsC_1_4_MM7 | Phospholipid/cardiolipin biosynthesis | Fatty Acid Metabolism | TCGTGGCCGCTCGCCAGCCG | 0 | 1 |
| dxs_1_17_C | Isoprenoid Biosynthesis | Ubiquinone Biosynthesis | CAGTGCCAGGGTCGGGTATT | 1 | 1 |
| dxs_1_17_MM7 | Isoprenoid Biosynthesis | Ubiquinone Biosynthesis | GTCACGGAGGGTCGGGTATT | 1 | 1 |
| umuC_1_35_C | SOS response | Stress Responses | GCGAAACACCGTCTCACAGC | 0 | 1 |

| | | | | | |
|---|---|---|---|---|---|
| umuC_1_35_MM7 | SOS response | Stress Responses | CGCTTTGACCGTCTCACAGC | 0 | 0 |
| aspC_1_17_C | Aspartate Biosynthesis | Amino Acid Biosynthesis | AATCGGGTCGGCAGGAGCGG | 0 | 0 |
| aspC_1_17_MM7 | Aspartate Biosynthesis | Amino Acid Biosynthesis | TTAGCCCTCGGCAGGAGCGG | 0 | 1 |
| dgkA_1_14_C | Phospholipid/cardiolipin biosynthesis | Fatty Acid Metabolism | GATAATTCGGGTGAATCCAG | 0 | 1 |
| dgkA_1_14_MM7 | Phospholipid/cardiolipin biosynthesis | Fatty Acid Metabolism | CTATTAACGGGTGAATCCAG | 0 | 0 |
| fadA_1_35_C | β-oxidation | Fatty Acid Metabolism | GCCCTTCGAACGGCCCATCG | 0 | 1 |
| fadA_1_35_MM7 | β-oxidation | Fatty Acid Metabolism | CGGGAAGGAACGGCCCATCG | 0 | 1 |
| poxB_1_29_C | Pyruvate metabolism | Central Carbon Metabolism | CCCTGCCGATTCGAGTGTTT | 0 | 1 |
| poxB_1_29_MM7 | Pyruvate metabolism | Central Carbon Metabolism | GGGACGGGATTCGAGTGTTT | 0 | 0 |
| fadR_1_21_C | Isoprenoid Biosynthesis | Ubiquinone Biosynthesis | ACTCTTCCGCGAAACCCGCC | 0 | 0 |
| fadR_1_21_MM7 | Isoprenoid Biosynthesis | Ubiquinone Biosynthesis | TGAGAAGCGCGAAACCCGCC | 0 | 0 |

| | | | | | |
|---|---|---|---|---|---|
| aroH_1_21_C | Chorismate Biosynthesis | Amino Acid Biosynthesis | GGCTCTCAAT ACGCGCAGTA | 0 | 1 |
| aroH_1_21_MM7 | Chorismate Biosynthesis | Amino Acid Biosynthesis | CCGAGAGAAT ACGCGCAGTA | 0 | 0 |
| pabA_1_35_C | Folate Biosynthesis | Folate Biosynthesis | AAAGTACTGG TAGAGGTTCC | 0 | 1 |
| pabA_1_35_MM7 | Folate Biosynthesis | Folate Biosynthesis | TTTCATGTGG TAGAGGTTCC | 0 | 1 |
| gdhA_1_42 | TCA cycle | Central Carbon Metabolism | TTTGATTCGG GTCGCGCTTT | 0 | 1 |
| uvrC_1_38_C | SOS response | Stress Responses | AACGCCTGGC TGGCTGGTTA | 0 | 0 |
| uvrC_1_38_MM7 | SOS response | Stress Responses | TTGCGGAGGC TGGCTGGTTA | 0 | 0 |
| recF_1_5_C | SOS response | Stress Responses | GCGGATCAAC AAGCGGGTGA | 0 | 1 |
| recF_1_5_MM7 | SOS response | Stress Responses | CGCCTAGAAC AAGCGGGTGA | 0 | 0 |
| fabI_1_14_C | Fatty Acid Biosynthesis | Fatty Acid Metabolism | GGTTACCAGA ATGCGCTTAC | 1 | 1 |
| fabI_1_14_MM7 | Fatty Acid Biosynthesis | Fatty Acid Metabolism | CCAATGGAGA ATGCGCTTAC | 1 | 1 |

| | | | | | |
|---|---|---|---|---|---|
| aceB_1_23_C | TCA cycle | Central Carbon Metabolism | CCTTGTGAAA GCCAGTTCAT | 0 | 1 |
| aceB_1_23_MM 7 | TCA cycle | Central Carbon Metabolism | GGAACACAAA GCCAGTTCAT | 0 | 0 |
| leuC_1_65_C | Branched Chain Amino Acid Metabolism | Amino Acid Biosynthesis | GCGGTCGATA TATAACAGTG | 0 | 1 |
| leuC_1_65_MM 7 | Branched Chain Amino Acid Metabolism | Amino Acid Biosynthesis | CGCCAGCATA TATAACAGTG | 0 | 1 |
| zwf_1_20_C | Pentose phosphate pathway | Pentose Phosphate Pathway | AATGACCAGG TCACAGGCCT | 0 | 0 |
| zwf_1_20_MM7 | Pentose phosphate pathway | Pentose Phosphate Pathway | TTACTGGAGG TCACAGGCCT | 0 | 0 |
| ppc_1_65_C | Gluconeogenesis | Central Carbon Metabolism | TTCTCCCAAC GCATCCTTGA | 0 | 0 |
| ppc_1_65_MM7 | Gluconeogenesis | Central Carbon Metabolism | AAGAGGGAAC GCATCCTTGA | 0 | 0 |
| fumA_1_66_C | TCA cycle | Central Carbon Metabolism | TAACGTGTTC GCTGGTTAGC | 0 | 0 |
| fumA_1_66_MM 7 | TCA cycle | Central Carbon Metabolism | ATTGCACTTC GCTGGTTAGC | 0 | 0 |
| ubiD_1_81_C | ubiquinone (coenzyme Q) biosynthesis | Ubiquinone Biosynthesis | TTTCCAGATG CGGATCCACC | 1 | 0 |

| | | | | | |
|---|---|---|---|---|---|
| ubiD_1_81_MM7 | ubiquinone (coenzyme Q) biosynthesis | Ubiquinone Biosynthesis | AAAGGTCATGCGGATCCACC | 1 | 0 |
| ilvI_1_20_C | Branched Chain Amino Acid Metabolism | Amino Acid Biosynthesis | AAGCGATCGGACGACCATCT | 0 | 1 |
| ilvI_1_20_MM7 | Branched Chain Amino Acid Metabolism | Amino Acid Biosynthesis | TTCGCTACGGACGACCATCT | 0 | 1 |
| pheA_1_15_C | Branched Chain Amino Acid Metabolism | Amino Acid Biosynthesis | TCTCTCGCAGCGCCAGTAAC | 0 | 0 |
| pheA_1_15_MM7 | Branched Chain Amino Acid Metabolism | Amino Acid Biosynthesis | AGAGAGCCAGCGCCAGTAAC | 0 | 0 |
| dmlA_1_30_C | Branched Chain Amino Acid Metabolism | Amino Acid Biosynthesis | CTTTGCCAATCCCGTCTCCC | 0 | 0 |
| dmlA_1_30_MM7 | Branched Chain Amino Acid Metabolism | Amino Acid Biosynthesis | GAAACGGAATCCCGTCTCCC | 0 | 0 |
| pykA_1_35_C | Glycolysis | Central Carbon Metabolism | ATCTGTTGCTGGGCCTAACG | 0 | 1 |
| pykA_1_35_MM7 | Glycolysis | Central Carbon Metabolism | TAGACAAGCTGGGCCTAACG | 0 | 1 |
| tyrA_1_127_C | Aromatic amino acid biosynthesis | Amino Acid Biosynthesis | TCGCGCTCCGGAACATAAAT | 0 | 1 |
| tyrA_1_127_MM7 | Aromatic amino acid biosynthesis | Amino Acid Biosynthesis | AGCGCGACCGGAACATAAAT | 0 | 1 |

| | | | | | |
|---|---|---|---|---|---|
| fadD_1_21_C | β-oxidation | Fatty Acid Metabolism | TCGGAACGTC CGCGGGATAA | 0 | 1 |
| fadD_1_21_MM 7 | β-oxidation | Fatty Acid Metabolism | AGCCTTGGTC CGCGGGATAA | 0 | 0 |
| aroE_1_28_C | Chorismate Biosynthesis | Amino Acid Biosynthesis | GGCGATTTGC TGTGGGCTAT | 0 | 0 |
| aroE_1_28_MM 7 | Chorismate Biosynthesis | Amino Acid Biosynthesis | CCGCTAATGC TGTGGGCTAT | 0 | 1 |
| maeB_1_36_C | Malate metabolism | Central Carbon Metabolism | TCCCTGGAAC TGGAAATTCA | 0 | 1 |
| maeB_1_36_M M7 | Malate metabolism | Central Carbon Metabolism | AGGGACCAAC TGGAAATTCA | 0 | 0 |
| accA_1_31_C | Fatty Acid Biosynthesis | Fatty Acid Metabolism | TTCGCTTCCA GCTCTGCAAT | 1 | 1 |
| accA_1_31_MM 7 | Fatty Acid Biosynthesis | Fatty Acid Metabolism | AAGCGAACCA GCTCTGCAAT | 1 | 1 |
| aroF_1_58_C | Chorismate Biosynthesis | Amino Acid Biosynthesis | AAAGCGGCCT TCAGTTGTTC | 0 | 0 |
| aroF_1_58_MM 7 | Chorismate Biosynthesis | Amino Acid Biosynthesis | TTTCGCCCCT TCAGTTGTTC | 0 | 1 |
| fabF_1_26_C | Fatty Acid Biosynthesis | Fatty Acid Metabolism | AGGAGACAAC ATGCCCAGTC | 0 | 1 |

| | | | | | |
|---|---|---|---|---|---|
| fabF_1_26_MM7 | Fatty Acid Biosynthesis | Fatty Acid Metabolism | TCCTCTGAAC ATGCCCAGTC | 0 | 0 |
| ridA_1_31_C | Branched Chain Amino Acid Metabolism | Amino Acid Biosynthesis | ACGTAAGGAC CGATAGCTGC | 0 | 0 |
| ridA_1_31_MM7 | Branched Chain Amino Acid Metabolism | Amino Acid Biosynthesis | TGCATTCGAC CGATAGCTGC | 0 | 1 |
| psd_1_40_C | Phospholipid/cardi olipin biosynthesis | Fatty Acid Metabolism | AGGCGAGTAA GCCATAGTTT | 1 | 1 |
| psd_1_40_MM7 | Phospholipid/cardi olipin biosynthesis | Fatty Acid Metabolism | TCCGCTCTAA GCCATAGTTT | 1 | 0 |
| fadE_1_33_C | β-oxidation | Fatty Acid Metabolism | GATAGAACAA CGCGCCGAG C | 0 | 0 |
| fadE_1_33_MM 7 | β-oxidation | Fatty Acid Metabolism | CTATCTTCAA CGCGCCGAG C | 0 | 1 |
| sdhB_1_33_C | TCA cycle | Central Carbon Metabolism | GCGGAGCATC ATCAACATCC | 0 | 1 |
| sdhB_1_33_MM 7 | TCA cycle | Central Carbon Metabolism | CGCCTCGATC ATCAACATCC | 0 | 0 |
| bacA_1_70_C | Isoprenoid Biosynthesis | Ubiquinone Biosynthesis | ATATGGCCCG TGCTGGATAC | 0 | 0 |
| bacA_1_70_MM 7 | Isoprenoid Biosynthesis | Ubiquinone Biosynthesis | TATACCGCCG TGCTGGATAC | 0 | 1 |

| | | | | | |
|---|---|---|---|---|---|
| mutL_1_4_C | SOS response | Stress Responses | TGTGGCGGTA AGACCTGAAT | 0 | 1 |
| mutL_1_4_MM7 | SOS response | Stress Responses | ACACCGCGTA AGACCTGAAT | 0 | 0 |
| ruvB_1_15_C | SOS response | Stress Responses | TGGTACCGGC AGAAATCAGA | 0 | 0 |
| ruvB_1_15_MM 7 | SOS response | Stress Responses | ACCATGGGGC AGAAATCAGA | 0 | 0 |
| mutS_1_26_C | SOS response | Stress Responses | CTGCTGCATC ATGGGCGTAT | 0 | 0 |
| mutS_1_26_MM 7 | SOS response | Stress Responses | GACGACGATC ATGGGCGTAT | 0 | 1 |
| gltA_1_26_C | TCA cycle | Central Carbon Metabolism | AACAGCTGTA TCCCCGTTGA | 0 | 0 |
| gltA_1_26_MM7 | TCA cycle | Central Carbon Metabolism | TTGTCGAGTA TCCCCGTTGA | 0 | 0 |
| gapA_1_54_C | Glycolysis | Central Carbon Metabolism | CAGAACGTTT CTGAGCAGCA | 1 | 1 |
| gapA_1_54_MM 7 | Glycolysis | Central Carbon Metabolism | GTCTTGCTTT CTGAGCAGCA | 1 | 0 |
| dinB_1_41_C | SOS response | Stress Responses | ATTGTCGCGC ATCTCCACTG | 0 | 1 |

| | | | | | |
|---|---|---|---|---|---|
| dinB_1_41_MM7 | SOS response | Stress Responses | TAACAGCCGC ATCTCCACTG | 0 | 0 |
| livK_1_86_C | Stringent response | Stress Responses | GCCGGACATC GCGCCGACAA | 0 | 1 |
| livK_1_86_MM7 | Stringent response | Stress Responses | CGGCCTGATC GCGCCGACAA | 0 | 1 |
| hpf_1_17_C | Stringent response | Stress Responses | GGTGATCTCG ACGTTATTTC | 0 | 1 |
| hpf_1_17_MM7 | Stringent response | Stress Responses | CCACTAGTCG ACGTTATTTC | 0 | 0 |
| pabC_1_80_C | Folate Biosynthesis | Folate Biosynthesis | ACCGTCGATA ACTCTGGCGG | 0 | 1 |
| pabC_1_80_MM7 | Folate Biosynthesis | Folate Biosynthesis | TGGCAGCATA ACTCTGGCGG | 0 | 1 |
| plsB_1_5_C | Phospholipid/cardiolipin biosynthesis | Fatty Acid Metabolism | GTAGTAAATT CGTGGCCAG C | 1 | 1 |
| plsB_1_5_MM7 | Phospholipid/cardiolipin biosynthesis | Fatty Acid Metabolism | CATCATTATTC GTGGCCAGC | 1 | 0 |
| yhbV_1_11_C | ubiquinone (coenzyme Q) biosynthesis | Ubiquinone Biosynthesis | GTACCACAGC ACTGGCCCTA | 0 | 0 |
| yhbV_1_11_MM7 | ubiquinone (coenzyme Q) biosynthesis | Ubiquinone Biosynthesis | CATGGTGAGC ACTGGCCCTA | 0 | 0 |

| | | | | | |
|---|---|---|---|---|---|
| eda_1_33_C | Entner-Doudoroff shunt | Central Carbon Metabolism | GTACAACCGG GCCGGTGGT C | 0 | 1 |
| eda_1_33_MM7 | Entner-Doudoroff shunt | Central Carbon Metabolism | CATGTTGCGG GCCGGTGGT C | 0 | 0 |
| ilvB_1_66_C | Branched Chain Amino Acid Metabolism | Amino Acid Biosynthesis | TCTTAATGCC CTGCTGTTCC | 0 | 1 |
| ilvB_1_66_MM7 | Branched Chain Amino Acid Metabolism | Amino Acid Biosynthesis | AGAATTAGCC CTGCTGTTCC | 0 | 0 |
| uvrD_1_15_C | SOS response | Stress Responses | TGTCATTAAG GCTGTCGAGC | 0 | 1 |
| uvrD_1_15_MM 7 | SOS response | Stress Responses | ACAGTAAAAG GCTGTCGAGC | 0 | 1 |
| recO_1_27_C | SOS response | Stress Responses | CGCTCCACGG GCGACTATGC | 0 | 1 |
| recO_1_27_MM 7 | SOS response | Stress Responses | GCGAGGTCG GGCGACTATG C | 0 | 1 |
| ruvA_1_40_C | SOS response | Stress Responses | ACTTCAATTAA CACCAGCGG | 0 | 1 |
| ruvA_1_40_MM 7 | SOS response | Stress Responses | TGAAGTTTTA ACACCAGCGG | 0 | 1 |
| leuD_1_30_C | Branched Chain Amino Acid Metabolism | Amino Acid Biosynthesis | CGGCATCCAG CGGAACCACC | 0 | 1 |

| | | | | | |
|---|---|---|---|---|---|
| leuD_1_30_MM7 | Branched Chain Amino Acid Metabolism | Amino Acid Biosynthesis | GCCGTAGCAGCGGAACCACC | 0 | 1 |
| lexA_1_17_C | SOS response | Stress Responses | ATCAAACACCTCTTGTTGCC | 1 | 0 |
| lexA_1_17_MM7 | SOS response | Stress Responses | TAGTTTGACCTCTTGTTGCC | 1 | 1 |
| sdhC_1_28_C | TCA cycle | Central Carbon Metabolism | GTCTGTAGGTCCAGATTAAC | 0 | 1 |
| sdhC_1_28_MM7 | TCA cycle | Central Carbon Metabolism | CAGACATGGTCCAGATTAAC | 0 | 1 |
| mdh_1_15_C | TCA cycle | Central Carbon Metabolism | CAATACCGCCAGCAGCGCCG | 0 | 0 |
| mdh_1_15_MM7 | TCA cycle | Central Carbon Metabolism | GTTATGGGCCAGCAGCGCCG | 0 | 0 |
| acnA_1_11_C | TCA cycle | Central Carbon Metabolism | GTCCTTACTGGCTTCTCGTA | 0 | 1 |
| acnA_1_11_MM7 | TCA cycle | Central Carbon Metabolism | CAGGAATCTGGCTTCTCGTA | 0 | 0 |
| rmf_1_24_C | Stringent response | Stress Responses | CACGTTGATGTGCCCGTTCC | 0 | 1 |
| rmf_1_24_MM7 | Stringent response | Stress Responses | GTGCAACATGTGCCCGTTCC | 0 | 0 |

| | | | | | |
|---|---|---|---|---|---|
| cdsA_1_40_C | Phospholipid/cardiolipin biosynthesis | Fatty Acid Metabolism | AACAACGCCGCGATGACGAC | 1 | 0 |
| cdsA_1_40_MM7 | Phospholipid/cardiolipin biosynthesis | Fatty Acid Metabolism | TTGTTGCCCGCGATGACGAC | 1 | 1 |
| pgpA_1_13_C | Phospholipid/cardiolipin biosynthesis | Fatty Acid Metabolism | TTCGCGACATCTTTATGGCG | 0 | 1 |
| pgpA_1_13_MM7 | Phospholipid/cardiolipin biosynthesis | Fatty Acid Metabolism | AAGCGCTCATCTTTATGGCG | 0 | 0 |
| ispB_1_29_C | Isoprenoid Biosynthesis | Ubiquinone Biosynthesis | AACACCCGCCATATCTTGCG | 1 | 1 |
| ispB_1_29_MM7 | Isoprenoid Biosynthesis | Ubiquinone Biosynthesis | TTGTGGGGCCATATCTTGCG | 1 | 0 |
| fadJ_1_50_C | β-oxidation | Fatty Acid Metabolism | CGGTACGTCGATGGTGATAA | 0 | 1 |
| fadJ_1_50_MM7 | β-oxidation | Fatty Acid Metabolism | GCCATGCTCGATGGTGATAA | 0 | 1 |
| pfkB_1_31_C | Glycolysis | Central Carbon Metabolism | ATTGTTGCGCTATCGAGAGA | 0 | 0 |
| pfkB_1_31_MM7 | Glycolysis | Central Carbon Metabolism | TAACAACCGCTATCGAGAGA | 0 | 0 |
| aceE_1_15_C | TCA cycle | Central Carbon Metabolism | CGATCGGATCCACGTCATTT | 0 | 0 |

| | | | | | |
|---|---|---|---|---|---|
| aceE_1_15_MM7 | TCA cycle | Central Carbon Metabolism | GCTAGCCATC CACGTCATTT | 0 | 1 |
| aroB_1_73_C | Chorismate Biosynthesis | Amino Acid Biosynthesis | TTCAGCGGTA AGAATGAAGC | 0 | 0 |
| aroB_1_73_MM7 | Chorismate Biosynthesis | Amino Acid Biosynthesis | AAGTCGCGTA AGAATGAAGC | 0 | 0 |
| fabA_1_36_C | Fatty Acid Biosynthesis | Fatty Acid Metabolism | CACCGCGACC AGAGGCAAGA | 1 | 1 |
| fabA_1_36_MM7 | Fatty Acid Biosynthesis | Fatty Acid Metabolism | GTGGCGCAC CAGAGGCAAG A | 1 | 1 |
| ybjG_1_49_C | Isoprenoid Biosynthesis | Ubiquinone Biosynthesis | ATCATCCACG GAGCCGAGTC | 0 | 1 |
| ybjG_1_49_MM7 | Isoprenoid Biosynthesis | Ubiquinone Biosynthesis | TAGTAGGACG GAGCCGAGTC | 0 | 0 |
| gnd_1_5_C | Pentose phosphate pathway | Pentose Phosphate Pathway | GACTACGCCG ATCTGTTGCT | 0 | 1 |
| gnd_1_5_MM7 | Pentose phosphate pathway | Pentose Phosphate Pathway | CTGATGCCCG ATCTGTTGCT | 0 | 0 |
| accD_1_37_C | Fatty Acid Biosynthesis | Fatty Acid Metabolism | GGAATGCTCG CCTTGCGGGT | 1 | 1 |
| accD_1_37_MM7 | Fatty Acid Biosynthesis | Fatty Acid Metabolism | CCTTACGTCG CCTTGCGGGT | 1 | 1 |

| | | | | | |
|---|---|---|---|---|---|
| fis_1_76_C | Stringent response | Stress Responses | TGTTTAACCG AGTCACGCAG | 0 | 0 |
| fis_1_76_MM7 | Stringent response | Stress Responses | ACAAATTCCG AGTCACGCAG | 0 | 1 |
| edd_1_101_C | Entner-Doudoroff shunt | Central Carbon Metabolism | TGCCAACTGC GAACGATGAA | 0 | 1 |
| edd_1_101_MM7 | Entner-Doudoroff shunt | Central Carbon Metabolism | ACGGTTGTGC GAACGATGAA | 0 | 0 |
| fabH_1_37_C | Fatty Acid Biosynthesis | Fatty Acid Metabolism | GCGTTTGTCC GCACTTGTTC | 1 | 0 |
| fabH_1_37_MM7 | Fatty Acid Biosynthesis | Fatty Acid Metabolism | CGCAAACTCC GCACTTGTTC | 1 | 0 |
| tpiA_1_45_C | Gluconeogenesis | Central Carbon Metabolism | CCAGCTCGTG AACCATGTGG | 0 | 1 |
| tpiA_1_45_MM7 | Gluconeogenesis | Central Carbon Metabolism | GGTCGAGGT GAACCATGTG G | 0 | 1 |
| hisG_1_41_C | Stringent response | Stress Responses | TGAGTCATCA CTTAAACGGC | 0 | 1 |
| hisG_1_41_MM7 | Stringent response | Stress Responses | ACTCAGTTCA CTTAAACGGC | 0 | 1 |
| plsC_1_75_C | Phospholipid/cardiolipin biosynthesis | Fatty Acid Metabolism | TCGGGTTACG CGGGCTGAAA | 1 | 0 |

| | | | | | |
|---|---|---|---|---|---|
| plsC_1_75_MM7 | Phospholipid/cardiolipin biosynthesis | Fatty Acid Metabolism | AGCCCAAACGCGGGCTGAAA | 1 | 1 |
| pflB_1_47_C | Fermentation | Central Carbon Metabolism | TTCATTCTGCCAGTCACCTT | 0 | 0 |
| pflB_1_47_MM7 | Fermentation | Central Carbon Metabolism | AAGTAAGTGCCAGTCACCTT | 0 | 1 |
| recA_1_77_C | SOS response | Stress Responses | GTCTTCACCCAGGCGCATGA | 0 | 0 |
| recA_1_77_MM7 | SOS response | Stress Responses | CAGAAGTCCCAGGCGCATGA | 0 | 0 |
| ispF_1_32_C | Isoprenoid Biosynthesis | Ubiquinone Biosynthesis | AATTGGGCCTTCACCGCCAA | 1 | 0 |
| ispF_1_32_MM7 | Isoprenoid Biosynthesis | Ubiquinone Biosynthesis | TTAACCCCCTTCACCGCCAA | 1 | 0 |
| sucD_1_26_C | TCA cycle | Central Carbon Metabolism | AAAGCCCTGGCAGATAACCT | 0 | 0 |
| sucD_1_26_MM7 | TCA cycle | Central Carbon Metabolism | TTTCGGGTGGCAGATAACCT | 0 | 1 |
| yhbU_1_15_C | ubiquinone (coenzyme Q) biosynthesis | Ubiquinone Biosynthesis | GCGCCGGGAGATTTCCGGCA | 0 | 0 |
| yhbU_1_15_MM7 | ubiquinone (coenzyme Q) biosynthesis | Ubiquinone Biosynthesis | CGCGGCCGAGATTTCCGGCA | 0 | 1 |

| | | | | | |
|---|---|---|---|---|---|
| rpiA_1_58_C | Pentose phosphate pathway | Pentose Phosphate Pathway | CCTACACCAA CAATGGTGCC | 0 | 0 |
| rpiA_1_58_MM7 | Pentose phosphate pathway | Pentose Phosphate Pathway | GGATGTGCAA CAATGGTGCC | 0 | 0 |
| tktB_1_15_C | Pentose phosphate pathway | Pentose Phosphate Pathway | GTGCGCGAAT CGCATTGGCA | 0 | 1 |
| tktB_1_15_MM7 | Pentose phosphate pathway | Pentose Phosphate Pathway | CACGCGCAAT CGCATTGGCA | 0 | 1 |
| spoT_1_38_C | Stringent response | Stress Responses | GATTTGGTCT TCCGGCAGGT | 0 | 1 |
| spoT_1_38_MM 7 | Stringent response | Stress Responses | CTAAACCTCT TCCGGCAGGT | 0 | 0 |
| ubiA_1_64_C | ubiquinone (coenzyme Q) biosynthesis | Ubiquinone Biosynthesis | AGCAGCAGTA ACGCGCCAAT | 1 | 1 |
| ubiA_1_64_MM7 | ubiquinone (coenzyme Q) biosynthesis | Ubiquinone Biosynthesis | TCGTCGTGTA ACGCGCCAAT | 1 | 1 |
| fadI_1_16_C | β-oxidation | Fatty Acid Metabolism | TCGCCCTGGC GGGTAACCAG | 0 | 0 |
| fadI_1_16_MM7 | β-oxidation | Fatty Acid Metabolism | AGCGGGAGG CGGGTAACCA G | 0 | 0 |
| clpP_1_34_C | Stringent response | Stress Responses | ATCGGCACCA GCGCCATATG | 0 | 1 |

| | | | | | |
|---|---|---|---|---|---|
| clpP_1_34_MM7 | Stringent response | Stress Responses | TAGCCGTCCA GCGCCATATG | 0 | 1 |
| livJ_1_104_C | Stringent response | Stress Responses | ACCGTACTGC GCAACCGGAC | 0 | 1 |
| livJ_1_104_MM7 | Stringent response | Stress Responses | TGGCATGTGC GCAACCGGAC | 0 | 1 |
| pgpB_1_14_C | Phospholipid/cardi olipin biosynthesis | Fatty Acid Metabolism | AGCTCCCACT GCGGTACGTC | 0 | 1 |
| pgpB_1_14_MM 7 | Phospholipid/cardi olipin biosynthesis | Fatty Acid Metabolism | TCGAGGGACT GCGGTACGTC | 0 | 0 |
| prpC_1_76_C | TCA cycle | Central Carbon Metabolism | CAGAGCGCC GTATTGCCCG C | 0 | 1 |
| prpC_1_76_MM 7 | TCA cycle | Central Carbon Metabolism | GTCTCGCCCG TATTGCCCGC | 0 | 0 |
| sucB_1_24_C | TCA cycle | Central Carbon Metabolism | CTACGGATTC AGGCAGGTCA | 1 | 1 |
| sucB_1_24_MM 7 | TCA cycle | Central Carbon Metabolism | GATGCCTTTC AGGCAGGTCA | 1 | 1 |
| leuB_1_15_C | Branched Chain Amino Acid Metabolism | Amino Acid Biosynthesis | CCCCCGGCAA TACGGCAATA | 0 | 1 |
| leuB_1_15_MM7 | Branched Chain Amino Acid Metabolism | Amino Acid Biosynthesis | GGGGGCCCA ATACGGCAAT A | 0 | 1 |

| | | | | | |
|---|---|---|---|---|---|
| idi_1_68_C | Isoprenoid Biosynthesis | Ubiquinone Biosynthesis | GCGGGTGTCT GCCGTGTGTG | 0 | 1 |
| idi_1_68_MM7 | Isoprenoid Biosynthesis | Ubiquinone Biosynthesis | CGCCCACTCT GCCGTGTGTG | 0 | 0 |
| ispH_1_9_C | Isoprenoid Biosynthesis | Ubiquinone Biosynthesis | AACCACGCGG GTTGGCCAAC | 1 | 0 |
| ispH_1_9_MM7 | Isoprenoid Biosynthesis | Ubiquinone Biosynthesis | TTGGTGCCGG GTTGGCCAAC | 1 | 0 |
| elbB_1_121_C | Isoprenoid Biosynthesis | Ubiquinone Biosynthesis | ACATCAACCT GCTGCTTATC | 0 | 1 |
| elbB_1_121_MM7 | Isoprenoid Biosynthesis | Ubiquinone Biosynthesis | TGTAGTTCCT GCTGCTTATC | 0 | 1 |
| lon_1_7_C | Stringent response | Stress Responses | TCAATGCGTT CAGAACGCTC | 0 | 0 |
| lon_1_7_MM7 | Stringent response | Stress Responses | AGTTACGGTT CAGAACGCTC | 0 | 0 |
| dinG_1_11_C | SOS response | Stress Responses | AATTTGCGCT TTAAGCGCGG | 0 | 0 |
| dinG_1_11_MM7 | SOS response | Stress Responses | TTAAACGGCT TTAAGCGCGG | 0 | 1 |
| argB_1_109_C | Stringent response | Stress Responses | CCGCCGTGCA CAATCACCAG | 0 | 1 |

| | | | | | |
|---|---|---|---|---|---|
| argB_1_109_MM7 | Stringent response | Stress Responses | GGCGGCAGCACAATCACCAG | 0 | 1 |
| pgpC_1_99_C | Phospholipid/cardiolipin biosynthesis | Fatty Acid Metabolism | CAAGTAACGCATTCAGCGGT | 0 | 0 |
| pgpC_1_99_MM7 | Phospholipid/cardiolipin biosynthesis | Fatty Acid Metabolism | GTTCATTCGCATTCAGCGGT | 0 | 0 |
| fbaA_1_28_C | Glycolysis/gluconeogensis | Central Carbon Metabolism | TCATCACCAGTGATTACGCC | 1 | 0 |
| fbaA_1_28_MM7 | Glycolysis/gluconeogensis | Central Carbon Metabolism | AGTAGTGCAGTGATTACGCC | 1 | 1 |
| talB_1_17_C | Pentose phosphate pathway | Pentose Phosphate Pathway | GGTGGTGTACTGACGAAGGG | 0 | 1 |
| talB_1_17_MM7 | Pentose phosphate pathway | Pentose Phosphate Pathway | CCACCACTACTGACGAAGGG | 0 | 1 |
| aroL_1_62_C | Chorismate Biosynthesis | Amino Acid Biosynthesis | ACGGTTAAGCGAATCGGCAA | 0 | 0 |
| aroL_1_62_MM7 | Chorismate Biosynthesis | Amino Acid Biosynthesis | TGCCAATAGCGAATCGGCAA | 0 | 1 |
| ilvC_1_33_C | Branched Chain Amino Acid Metabolism | Amino Acid Biosynthesis | TGCCCAGCTGTGCCAGCTGC | 0 | 1 |
| ilvC_1_33_MM7 | Branched Chain Amino Acid Metabolism | Amino Acid Biosynthesis | ACGGGTCCTGTGCCAGCTGC | 0 | 1 |

| | | | | | |
|---|---|---|---|---|---|
| tktA_1_5_C | Pentose phosphate pathway | Pentose Phosphate Pathway | ATTGGCAAGC TCTTTACGTG | 1 | 0 |
| tktA_1_5_MM7 | Pentose phosphate pathway | Pentose Phosphate Pathway | TAACCGTAGC TCTTTACGTG | 1 | 0 |
| leuA_1_29_C | Branched Chain Amino Acid Metabolism | Amino Acid Biosynthesis | CTGTTCACCG TCGCGCAATG | 0 | 0 |
| leuA_1_29_MM7 | Branched Chain Amino Acid Metabolism | Amino Acid Biosynthesis | GACAAGTCCG TCGCGCAATG | 0 | 1 |
| uvrB_1_37_C | SOS response | Stress Responses | GCCTCTGGCT GATCGCCAGA | 0 | 1 |
| uvrB_1_37_MM 7 | SOS response | Stress Responses | CGGAGACGCT GATCGCCAGA | 0 | 1 |
| ruvC_1_25_C | SOS response | Stress Responses | TAGCCGGTCA CGCGCGAAC C | 0 | 1 |
| ruvC_1_25_MM 7 | SOS response | Stress Responses | ATCGGCCTCA CGCGCGAAC C | 0 | 1 |
| fabR_1_32_C | Fatty Acid Biosynthesis | Fatty Acid Metabolism | GGCTTCCACC AGCGAACGG C | 0 | 1 |
| fabR_1_32_MM 7 | Fatty Acid Biosynthesis | Fatty Acid Metabolism | CCGAAGGACC AGCGAACGG C | 0 | 1 |
| yqeF_1_37_C | Acetate Metabolism | Central Carbon Metabolism | GCACCACGAA AGCAGCCGAT | 0 | 0 |

| | | | | | |
|---|---|---|---|---|---|
| yqeF_1_37_MM7 | Acetate Metabolism | Central Carbon Metabolism | CGTGGTGGAAAGCAGCCGAT | 0 | 0 |
| ispE_1_53_C | Isoprenoid Biosynthesis | Ubiquinone Biosynthesis | GTAACCATCCGCACGCTGAC | 1 | 1 |
| ispE_1_53_MM7 | Isoprenoid Biosynthesis | Ubiquinone Biosynthesis | CATTGGTTCCGCACGCTGAC | 1 | 0 |
| clsB_1_64_C | Phospholipid/cardiolipin biosynthesis | Fatty Acid Metabolism | CCAATCGCCTTAAACACCGC | 0 | 0 |
| clsB_1_64_MM7 | Phospholipid/cardiolipin biosynthesis | Fatty Acid Metabolism | GGTTAGCCCTTAAACACCGC | 0 | 1 |
| pta_1_24_C | Acetate metabolism | Central Carbon Metabolism | GACCGACGCTGGTTCCGGTA | 0 | 1 |
| pta_1_24_MM7 | Acetate metabolism | Central Carbon Metabolism | CTGGCTGGCTGGTTCCGGTA | 0 | 1 |
| fabB_1_24_C | Fatty Acid Biosynthesis | Fatty Acid Metabolism | CGATGCTGGAAACAATGCCC | 1 | 0 |
| fabB_1_24_MM7 | Fatty Acid Biosynthesis | Fatty Acid Metabolism | GCTACGAGGAAACAATGCCC | 1 | 1 |

**Table S3:** Large-scale sgRNA library targeting *E. coli* genes throughout metabolism. Annotations of target gene cell process, pathway, and essentiality (Related to Figure 4). The last column indicates whether or not the sgRNA was recovered (identified by NGS) during our experimental screen.

| Guide | Replicates | OD600_avg | OD600_stdev | Lycopene (mg/(OD600*L)) avg | Lycopene (mg/(OD600*L) stdev |
|---|---|---|---|---|---|
| negC_rand_42 | 3 | 5.50 | 0.36 | 0.570 | 0.015 |
| acnB_1_15_C | 3 | 3.79 | 0.12 | 0.461 | 0.033 |
| acnB_1_15_MM7 | 3 | 5.32 | 0.27 | 0.591 | 0.077 |
| dxr_1_14_C | 0 | N.D. | N.D. | N.D. | N.D. |
| dxr_1_14_MM7 | 3 | 4.75 | 0.52 | 0.106 | 0.049 |
| fadB_1_20_C | 0 | N.D. | N.D. | N.D. | N.D. |
| fadB_1_20_MM7 | 0 | N.D. | N.D. | N.D. | N.D. |
| dnaA_1_32_C | 0 | N.D. | N.D. | N.D. | N.D. |
| dnaA_1_32_MM7 | 0 | N.D. | N.D. | N.D. | N.D. |
| dksA_1_32_C | 3 | 5.43 | 0.30 | 0.384 | 0.060 |
| dksA_1_32_MM7 | 3 | 5.56 | 0.35 | 0.396 | 0.039 |
| ispD_1_8_C | 3 | 5.79 | 0.74 | 0.229 | 0.023 |
| ispD_1_8_MM7 | 3 | 4.97 | 0.26 | 0.279 | 0.050 |
| fadH_1_9_C | 0 | N.D. | N.D. | N.D. | N.D. |
| fadH_1_9_MM7 | 3 | 5.87 | 0.42 | 0.652 | 0.031 |
| ispA_1_10_C | 3 | 6.50 | 0.21 | 0.293 | 0.032 |
| ispA_1_10_MM7 | 3 | 6.25 | 0.59 | 0.501 | 0.009 |
| sdhD_1_14_C | 0 | N.D. | N.D. | N.D. | N.D. |
| sdhD_1_14_MM7 | 3 | 6.58 | 0.41 | 0.698 | 0.032 |
| folP_1_14_C | 3 | 6.15 | 0.31 | 0.646 | 0.046 |
| folP_1_14_MM7 | 0 | N.D. | N.D. | N.D. | N.D. |
| aroD_1_83_C | 3 | 5.30 | 0.20 | 0.600 | 0.055 |
| aroD_1_83_MM7 | 0 | N.D. | N.D. | N.D. | N.D. |
| ubiG_1_27_C | 0 | N.D. | N.D. | N.D. | N.D. |
| ubiG_1_27_MM7 | 0 | N.D. | N.D. | N.D. | N.D. |
| trpE_1_16_C | 0 | N.D. | N.D. | N.D. | N.D. |
| trpE_1_16_MM7 | 0 | N.D. | N.D. | N.D. | N.D. |
| ilvA_1_16_C | 3 | 6.06 | 0.21 | 0.659 | 0.028 |
| ilvA_1_16_MM7 | 0 | N.D. | N.D. | N.D. | N.D. |
| sucA_1_23_C | 0 | N.D. | N.D. | N.D. | N.D. |
| sucA_1_23_MM7 | 0 | N.D. | N.D. | N.D. | N.D. |
| maeA_1_39_C | 0 | N.D. | N.D. | N.D. | N.D. |
| maeA_1_39_MM7 | 0 | N.D. | N.D. | N.D. | N.D. |
| yajO_1_26_C | 3 | 5.75 | 0.24 | 0.679 | 0.035 |

| | | | | | |
|---|---|---|---|---|---|
| yajO_1_26_MM7 | 3 | 6.20 | 0.96 | 0.661 | 0.128 |
| aroA_1_8_C | 3 | 6.25 | 0.41 | 0.508 | 0.034 |
| aroA_1_8_MM7 | 0 | N.D. | N.D. | N.D. | N.D. |
| polA_1_76_C | 3 | 5.88 | 0.21 | 0.621 | 0.075 |
| polA_1_76_MM7 | 3 | 5.53 | 0.21 | 0.583 | 0.021 |
| pck_1_23_C | 0 | N.D. | N.D. | N.D. | N.D. |
| pck_1_23_MM7 | 3 | 4.46 | 0.27 | 0.578 | 0.058 |
| aceF_1_22_C | 0 | N.D. | N.D. | N.D. | N.D. |
| aceF_1_22_MM7 | 3 | 5.38 | 0.12 | 0.768 | 0.052 |
| fbaB_1_70_C | 3 | 6.05 | 0.62 | 0.618 | 0.025 |
| fbaB_1_70_MM7 | 0 | N.D. | N.D. | N.D. | N.D. |
| gpsA_1_38_C | 3 | 2.93 | 0.82 | 0.528 | 0.157 |
| gpsA_1_38_MM7 | 0 | N.D. | N.D. | N.D. | N.D. |
| fumB_1_14_C | 0 | N.D. | N.D. | N.D. | N.D. |
| fumB_1_14_MM7 | 0 | N.D. | N.D. | N.D. | N.D. |
| icd_1_22_C | 0 | N.D. | N.D. | N.D. | N.D. |
| icd_1_22_MM7 | 3 | 5.38 | 0.14 | 0.528 | 0.048 |
| adhE_1_56_C | 3 | 5.61 | 0.28 | 0.600 | 0.063 |
| adhE_1_56_MM7 | 3 | 5.82 | 0.61 | 0.671 | 0.074 |
| pabB_1_68_C | 3 | 5.62 | 0.11 | 0.546 | 0.035 |
| pabB_1_68_MM7 | 0 | N.D. | N.D. | N.D. | N.D. |
| mutH_1_5_C | 0 | N.D. | N.D. | N.D. | N.D. |
| mutH_1_5_MM7 | 0 | N.D. | N.D. | N.D. | N.D. |
| fadL_1_15_C | 0 | N.D. | N.D. | N.D. | N.D. |
| fadL_1_15_MM7 | 0 | N.D. | N.D. | N.D. | N.D. |
| rpoD_1_12_C | 0 | N.D. | N.D. | N.D. | N.D. |
| rpoD_1_12_MM7 | 0 | N.D. | N.D. | N.D. | N.D. |
| pykF_1_179_C | 0 | N.D. | N.D. | N.D. | N.D. |
| pykF_1_179_MM7 | 0 | N.D. | N.D. | N.D. | N.D. |
| sdhA_1_44_C | 3 | 7.95 | 0.11 | 0.464 | 0.040 |
| sdhA_1_44_MM7 | 0 | N.D. | N.D. | N.D. | N.D. |
| aroK_1_128_C | 3 | 5.89 | 0.30 | 0.619 | 0.123 |
| aroK_1_128_MM7 | 3 | 5.65 | 0.27 | 0.662 | 0.030 |
| rpoS_1_86_C | 0 | N.D. | N.D. | N.D. | N.D. |
| rpoS_1_86_MM7 | 3 | 5.00 | 0.40 | 0.455 | 0.035 |
| menA_1_21_C | 0 | N.D. | N.D. | N.D. | N.D. |
| menA_1_21_MM7 | 0 | N.D. | N.D. | N.D. | N.D. |
| ilvH_1_71_C | 0 | N.D. | N.D. | N.D. | N.D. |
| ilvH_1_71_MM7 | 0 | N.D. | N.D. | N.D. | N.D. |
| gpmA_1_60_C | 3 | 5.96 | 0.47 | 0.647 | 0.027 |
| gpmA_1_60_MM7 | 0 | N.D. | N.D. | N.D. | N.D. |

| | | | | | |
|---|---|---|---|---|---|
| gpp_1_8_C | 3 | 5.85 | 0.14 | 0.575 | 0.006 |
| gpp_1_8_MM7 | 3 | 6.09 | 0.24 | 0.569 | 0.080 |
| accB_1_99_C | 3 | 2.40 | 0.76 | 0.882 | 0.176 |
| accB_1_99_MM7 | 3 | 2.33 | 0.52 | 0.894 | 0.203 |
| talA_1_32_C | 0 | N.D. | N.D. | N.D. | N.D. |
| talA_1_32_MM7 | 3 | 6.13 | 0.30 | 0.594 | 0.024 |
| ytjC_1_15_C | 3 | 5.28 | 0.33 | 0.638 | 0.060 |
| ytjC_1_15_MM7 | 3 | 5.75 | 0.17 | 0.682 | 0.041 |
| aceA_1_52_C | 3 | 5.77 | 0.49 | 0.576 | 0.054 |
| aceA_1_52_MM7 | 0 | N.D. | N.D. | N.D. | N.D. |
| sulA_1_89_C | 3 | 6.18 | 0.70 | 0.645 | 0.137 |
| sulA_1_89_MM7 | 0 | N.D. | N.D. | N.D. | N.D. |
| clsA_1_35_C | 0 | N.D. | N.D. | N.D. | N.D. |
| clsA_1_35_MM7 | 3 | 5.62 | 0.65 | 0.663 | 0.054 |
| ispU_1_40_C | 3 | 3.08 | 1.16 | 0.394 | 0.182 |
| ispU_1_40_MM7 | 0 | N.D. | N.D. | N.D. | N.D. |
| uvrA_1_54_C | 0 | N.D. | N.D. | N.D. | N.D. |
| uvrA_1_54_MM7 | 3 | 6.19 | 0.25 | 0.577 | 0.040 |
| lpd_1_45_C | 3 | 5.19 | 0.34 | 0.505 | 0.139 |
| lpd_1_45_MM7 | 3 | 4.57 | 0.12 | 0.448 | 0.127 |
| pgsA_1_15_C | 3 | 6.14 | 0.38 | 0.617 | 0.062 |
| pgsA_1_15_MM7 | 3 | 5.20 | 0.33 | 0.790 | 0.040 |
| rpe_1_20_C | 0 | N.D. | N.D. | N.D. | N.D. |
| rpe_1_20_MM7 | 3 | 5.99 | 0.07 | 0.535 | 0.038 |
| ldhA_1_45_C | 3 | 6.26 | 0.19 | 0.670 | 0.051 |
| ldhA_1_45_MM7 | 0 | N.D. | N.D. | N.D. | N.D. |
| entA_1_32_C | 3 | 5.64 | 0.20 | 0.630 | 0.030 |
| entA_1_32_MM7 | 3 | 5.72 | 0.28 | 0.632 | 0.047 |
| ispG_1_9_C | 0 | N.D. | N.D. | N.D. | N.D. |
| ispG_1_9_MM7 | 3 | 5.85 | 0.31 | 0.262 | 0.023 |
| pgi_1_16_C | 0 | N.D. | N.D. | N.D. | N.D. |
| pgi_1_16_MM7 | 3 | 5.78 | 0.33 | 0.601 | 0.036 |
| tyrB_1_20_C | 0 | N.D. | N.D. | N.D. | N.D. |
| tyrB_1_20_MM7 | 0 | N.D. | N.D. | N.D. | N.D. |
| aroC_1_38_C | 0 | N.D. | N.D. | N.D. | N.D. |
| aroC_1_38_MM7 | 3 | 4.96 | 0.42 | 0.707 | 0.061 |
| ilvM_1_87_C | 0 | N.D. | N.D. | N.D. | N.D. |
| ilvM_1_87_MM7 | 0 | N.D. | N.D. | N.D. | N.D. |
| ilvN_1_41_C | 0 | N.D. | N.D. | N.D. | N.D. |
| ilvN_1_41_MM7 | 0 | N.D. | N.D. | N.D. | N.D. |
| rpiB_1_104_C | 3 | 5.58 | 0.25 | 0.613 | 0.100 |

| | | | | | |
|---|---|---|---|---|---|
| rpiB_1_104_MM7 | 3 | 6.35 | 0.77 | 0.585 | 0.083 |
| pssA_1_39_C | 3 | 5.43 | 0.71 | 0.675 | 0.110 |
| pssA_1_39_MM7 | 3 | 5.80 | 1.04 | 0.619 | 0.081 |
| ptsG_1_55_C | 0 | N.D. | N.D. | N.D. | N.D. |
| ptsG_1_55_MM7 | 0 | N.D. | N.D. | N.D. | N.D. |
| ilvD_1_4_C | 3 | 5.81 | 0.45 | 0.724 | 0.065 |
| ilvD_1_4_MM7 | 0 | N.D. | N.D. | N.D. | N.D. |
| sucC_1_38_C | 3 | 6.67 | 0.70 | 0.665 | 0.087 |
| sucC_1_38_MM7 | 0 | N.D. | N.D. | N.D. | N.D. |
| accC_1_23_C | 0 | N.D. | N.D. | N.D. | N.D. |
| accC_1_23_MM7 | 0 | N.D. | N.D. | N.D. | N.D. |
| recR_1_8_C | 0 | N.D. | N.D. | N.D. | N.D. |
| recR_1_8_MM7 | 0 | N.D. | N.D. | N.D. | N.D. |
| relA_1_80_C | 3 | 5.54 | 0.23 | 0.591 | 0.025 |
| relA_1_80_MM7 | 0 | N.D. | N.D. | N.D. | N.D. |
| fumC_1_48_C | 3 | 5.64 | 0.16 | 0.695 | 0.075 |
| fumC_1_48_MM7 | 0 | N.D. | N.D. | N.D. | N.D. |
| pgk_1_20_C | 0 | N.D. | N.D. | N.D. | N.D. |
| pgk_1_20_MM7 | 3 | 6.25 | 0.49 | 0.751 | 0.070 |
| ilvE_1_120_C | 3 | 6.42 | 0.43 | 0.489 | 0.058 |
| ilvE_1_120_MM7 | 0 | N.D. | N.D. | N.D. | N.D. |
| clsC_1_4_C | 3 | 6.14 | 0.58 | 0.639 | 0.119 |
| clsC_1_4_MM7 | 3 | 6.46 | 0.38 | 0.615 | 0.076 |
| dxs_1_17_C | 3 | 4.96 | 0.17 | 0.054 | 0.009 |
| dxs_1_17_MM7 | 3 | 5.08 | 0.52 | 0.047 | 0.008 |
| umuC_1_35_C | 3 | 5.53 | 0.05 | 0.637 | 0.018 |
| umuC_1_35_MM7 | 0 | N.D. | N.D. | N.D. | N.D. |
| aspC_1_17_C | 0 | N.D. | N.D. | N.D. | N.D. |
| aspC_1_17_MM7 | 3 | 6.20 | 1.03 | 0.480 | 0.074 |
| dgkA_1_14_C | 3 | 5.55 | 0.00 | 0.773 | 0.061 |
| dgkA_1_14_MM7 | 0 | N.D. | N.D. | N.D. | N.D. |
| fadA_1_35_C | 3 | 6.19 | 0.64 | 0.612 | 0.055 |
| fadA_1_35_MM7 | 3 | 5.42 | 0.34 | 0.655 | 0.032 |
| poxB_1_29_C | 3 | 5.84 | 0.33 | 0.596 | 0.055 |
| poxB_1_29_MM7 | 0 | N.D. | N.D. | N.D. | N.D. |
| fadR_1_21_C | 0 | N.D. | N.D. | N.D. | N.D. |
| fadR_1_21_MM7 | 0 | N.D. | N.D. | N.D. | N.D. |
| aroH_1_21_C | 3 | 5.98 | 0.68 | 0.682 | 0.077 |
| aroH_1_21_MM7 | 0 | N.D. | N.D. | N.D. | N.D. |
| pabA_1_35_C | 3 | 7.39 | 0.43 | 0.594 | 0.037 |
| pabA_1_35_MM7 | 3 | 6.84 | 0.60 | 0.568 | 0.054 |

| | | | | | |
|---|---|---|---|---|---|
| gdhA_1_42 | 3 | 5.81 | 0.78 | 0.649 | 0.049 |
| uvrC_1_38_C | 0 | N.D. | N.D. | N.D. | N.D. |
| uvrC_1_38_MM7 | 0 | N.D. | N.D. | N.D. | N.D. |
| recF_1_5_C | 3 | 5.48 | 0.15 | 0.600 | 0.059 |
| recF_1_5_MM7 | 0 | N.D. | N.D. | N.D. | N.D. |
| fabI_1_14_C | 3 | 5.22 | 1.45 | 0.651 | 0.059 |
| fabI_1_14_MM7 | 3 | 5.76 | 0.48 | 0.581 | 0.043 |
| aceB_1_23_C | 3 | 5.80 | 0.40 | 0.631 | 0.021 |
| aceB_1_23_MM7 | 0 | N.D. | N.D. | N.D. | N.D. |
| leuC_1_65_C | 3 | 6.02 | 0.30 | 0.690 | 0.063 |
| leuC_1_65_MM7 | 3 | 6.45 | 0.56 | 0.680 | 0.033 |
| zwf_1_20_C | 0 | N.D. | N.D. | N.D. | N.D. |
| zwf_1_20_MM7 | 0 | N.D. | N.D. | N.D. | N.D. |
| ppc_1_65_C | 0 | N.D. | N.D. | N.D. | N.D. |
| ppc_1_65_MM7 | 0 | N.D. | N.D. | N.D. | N.D. |
| fumA_1_66_C | 0 | N.D. | N.D. | N.D. | N.D. |
| fumA_1_66_MM7 | 0 | N.D. | N.D. | N.D. | N.D. |
| ubiD_1_81_C | 0 | N.D. | N.D. | N.D. | N.D. |
| ubiD_1_81_MM7 | 0 | N.D. | N.D. | N.D. | N.D. |
| ilvI_1_20_C | 3 | 3.93 | 0.39 | 0.715 | 0.109 |
| ilvI_1_20_MM7 | 3 | 4.42 | 0.14 | 0.741 | 0.082 |
| pheA_1_15_C | 0 | N.D. | N.D. | N.D. | N.D. |
| pheA_1_15_MM7 | 0 | N.D. | N.D. | N.D. | N.D. |
| dmlA_1_30_C | 0 | N.D. | N.D. | N.D. | N.D. |
| dmlA_1_30_MM7 | 0 | N.D. | N.D. | N.D. | N.D. |
| pykA_1_35_C | 3 | 6.17 | 0.34 | 0.531 | 0.034 |
| pykA_1_35_MM7 | 3 | 5.75 | 0.07 | 0.563 | 0.054 |
| tyrA_1_127_C | 3 | 6.47 | 0.30 | 0.594 | 0.098 |
| tyrA_1_127_MM7 | 3 | 6.11 | 0.51 | 0.609 | 0.020 |
| fadD_1_21_C | 3 | 5.67 | 0.25 | 0.585 | 0.012 |
| fadD_1_21_MM7 | 0 | N.D. | N.D. | N.D. | N.D. |
| aroE_1_28_C | 0 | N.D. | N.D. | N.D. | N.D. |
| aroE_1_28_MM7 | 3 | 5.77 | 0.54 | 0.601 | 0.046 |
| maeB_1_36_C | 3 | 5.73 | 0.20 | 0.597 | 0.075 |
| maeB_1_36_MM7 | 0 | N.D. | N.D. | N.D. | N.D. |
| accA_1_31_C | 3 | 2.22 | 0.19 | 0.758 | 0.063 |
| accA_1_31_MM7 | 3 | 5.72 | 0.25 | 0.664 | 0.061 |
| aroF_1_58_C | 0 | N.D. | N.D. | N.D. | N.D. |
| aroF_1_58_MM7 | 3 | 5.80 | 0.18 | 0.656 | 0.011 |
| fabF_1_26_C | 3 | 5.36 | 0.57 | 0.500 | 0.048 |
| fabF_1_26_MM7 | 0 | N.D. | N.D. | N.D. | N.D. |

| | | | | | |
|---|---|---|---|---|---|
| ridA_1_31_C | 0 | N.D. | N.D. | N.D. | N.D. |
| ridA_1_31_MM7 | 3 | 5.74 | 0.47 | 0.609 | 0.011 |
| psd_1_40_C | 3 | 5.01 | 0.12 | 0.532 | 0.060 |
| psd_1_40_MM7 | 0 | N.D. | N.D. | N.D. | N.D. |
| fadE_1_33_C | 0 | N.D. | N.D. | N.D. | N.D. |
| fadE_1_33_MM7 | 3 | 6.01 | 0.39 | 0.574 | 0.058 |
| sdhB_1_33_C | 3 | 6.43 | 0.50 | 0.452 | 0.018 |
| sdhB_1_33_MM7 | 0 | N.D. | N.D. | N.D. | N.D. |
| bacA_1_70_C | 0 | N.D. | N.D. | N.D. | N.D. |
| bacA_1_70_MM7 | 3 | 5.61 | 0.46 | 0.640 | 0.069 |
| mutL_1_4_C | 3 | 6.20 | 0.52 | 0.594 | 0.047 |
| mutL_1_4_MM7 | 0 | N.D. | N.D. | N.D. | N.D. |
| ruvB_1_15_C | 0 | N.D. | N.D. | N.D. | N.D. |
| ruvB_1_15_MM7 | 0 | N.D. | N.D. | N.D. | N.D. |
| mutS_1_26_C | 0 | N.D. | N.D. | N.D. | N.D. |
| mutS_1_26_MM7 | 3 | 6.10 | 0.35 | 0.573 | 0.021 |
| gltA_1_26_C | 0 | N.D. | N.D. | N.D. | N.D. |
| gltA_1_26_MM7 | 0 | N.D. | N.D. | N.D. | N.D. |
| gapA_1_54_C | 3 | 5.70 | 0.68 | 0.678 | 0.113 |
| gapA_1_54_MM7 | 0 | N.D. | N.D. | N.D. | N.D. |
| dinB_1_41_C | 3 | 6.12 | 0.40 | 0.618 | 0.047 |
| dinB_1_41_MM7 | 0 | N.D. | N.D. | N.D. | N.D. |
| livK_1_86_C | 3 | 5.63 | 0.30 | 0.631 | 0.057 |
| livK_1_86_MM7 | 3 | 5.81 | 0.22 | 0.661 | 0.096 |
| hpf_1_17_C | 3 | 5.46 | 0.35 | 0.665 | 0.066 |
| hpf_1_17_MM7 | 0 | N.D. | N.D. | N.D. | N.D. |
| pabC_1_80_C | 3 | 5.78 | 0.49 | 0.609 | 0.102 |
| pabC_1_80_MM7 | 3 | 6.01 | 0.43 | 0.529 | 0.012 |
| plsB_1_5_C | 3 | 2.50 | 0.57 | 0.508 | 0.262 |
| plsB_1_5_MM7 | 0 | N.D. | N.D. | N.D. | N.D. |
| yhbV_1_11_C | 0 | N.D. | N.D. | N.D. | N.D. |
| yhbV_1_11_MM7 | 0 | N.D. | N.D. | N.D. | N.D. |
| eda_1_33_C | 3 | 5.93 | 0.33 | 0.596 | 0.044 |
| eda_1_33_MM7 | 0 | N.D. | N.D. | N.D. | N.D. |
| ilvB_1_66_C | 3 | 5.95 | 0.11 | 0.559 | 0.022 |
| ilvB_1_66_MM7 | 0 | N.D. | N.D. | N.D. | N.D. |
| uvrD_1_15_C | 3 | 6.50 | 0.38 | 0.408 | 0.023 |
| uvrD_1_15_MM7 | 3 | 5.71 | 0.20 | 0.673 | 0.025 |
| recO_1_27_C | 3 | 5.66 | 0.29 | 0.633 | 0.024 |
| recO_1_27_MM7 | 3 | 5.91 | 0.16 | 0.657 | 0.049 |
| ruvA_1_40_C | 3 | 5.19 | 0.50 | 0.732 | 0.068 |

| | | | | | |
|---|---|---|---|---|---|
| ruvA_1_40_MM7 | 3 | 5.98 | 0.26 | 0.619 | 0.032 |
| leuD_1_30_C | 3 | 5.69 | 0.45 | 0.564 | 0.064 |
| leuD_1_30_MM7 | 3 | 5.55 | 0.51 | 0.650 | 0.062 |
| lexA_1_17_C | 0 | N.D. | N.D. | N.D. | N.D. |
| lexA_1_17_MM7 | 3 | 6.14 | 0.34 | 0.617 | 0.111 |
| sdhC_1_28_C | 3 | 3.36 | 0.68 | 0.203 | 0.036 |
| sdhC_1_28_MM7 | 3 | 3.14 | 0.19 | 0.178 | 0.031 |
| mdh_1_15_C | 0 | N.D. | N.D. | N.D. | N.D. |
| mdh_1_15_MM7 | 0 | N.D. | N.D. | N.D. | N.D. |
| acnA_1_11_C | 3 | 5.89 | 0.20 | 0.714 | 0.018 |
| acnA_1_11_MM7 | 0 | N.D. | N.D. | N.D. | N.D. |
| rmf_1_24_C | 3 | 5.75 | 0.33 | 0.633 | 0.033 |
| rmf_1_24_MM7 | 0 | N.D. | N.D. | N.D. | N.D. |
| cdsA_1_40_C | 0 | N.D. | N.D. | N.D. | N.D. |
| cdsA_1_40_MM7 | 3 | 5.60 | 0.40 | 0.614 | 0.059 |
| pgpA_1_13_C | 3 | 5.99 | 0.21 | 0.642 | 0.064 |
| pgpA_1_13_MM7 | 0 | N.D. | N.D. | N.D. | N.D. |
| ispB_1_29_C | 3 | 4.67 | 1.08 | 0.532 | 0.205 |
| ispB_1_29_MM7 | 0 | N.D. | N.D. | N.D. | N.D. |
| fadJ_1_50_C | 3 | 5.61 | 0.27 | 0.587 | 0.023 |
| fadJ_1_50_MM7 | 3 | 5.46 | 0.29 | 0.582 | 0.034 |
| pfkB_1_31_C | 0 | N.D. | N.D. | N.D. | N.D. |
| pfkB_1_31_MM7 | 0 | N.D. | N.D. | N.D. | N.D. |
| aceE_1_15_C | 0 | N.D. | N.D. | N.D. | N.D. |
| aceE_1_15_MM7 | 3 | 5.94 | 0.20 | 0.697 | 0.024 |
| aroB_1_73_C | 0 | N.D. | N.D. | N.D. | N.D. |
| aroB_1_73_MM7 | 0 | N.D. | N.D. | N.D. | N.D. |
| fabA_1_36_C | 3 | 4.08 | 1.30 | 0.555 | 0.100 |
| fabA_1_36_MM7 | 3 | 5.29 | 1.23 | 0.529 | 0.044 |
| ybjG_1_49_C | 3 | 5.88 | 0.21 | 0.538 | 0.101 |
| ybjG_1_49_MM7 | 0 | N.D. | N.D. | N.D. | N.D. |
| gnd_1_5_C | 3 | 6.09 | 0.69 | 0.599 | 0.026 |
| gnd_1_5_MM7 | 0 | N.D. | N.D. | N.D. | N.D. |
| accD_1_37_C | 3 | 2.56 | 0.49 | 0.891 | 0.151 |
| accD_1_37_MM7 | 3 | 2.73 | 0.76 | 0.892 | 0.061 |
| fis_1_76_C | 0 | N.D. | N.D. | N.D. | N.D. |
| fis_1_76_MM7 | 3 | 6.32 | 0.57 | 0.573 | 0.039 |
| edd_1_101_C | 3 | 5.51 | 0.10 | 0.650 | 0.073 |
| edd_1_101_MM7 | 0 | N.D. | N.D. | N.D. | N.D. |
| fabH_1_37_C | 0 | N.D. | N.D. | N.D. | N.D. |
| fabH_1_37_MM7 | 0 | N.D. | N.D. | N.D. | N.D. |

| | | | | | |
|---|---|---|---|---|---|
| tpiA_1_45_C | 3 | 6.39 | 1.54 | 0.473 | 0.051 |
| tpiA_1_45_MM7 | 3 | 6.32 | 0.61 | 0.428 | 0.083 |
| hisG_1_41_C | 3 | 5.89 | 0.22 | 0.600 | 0.057 |
| hisG_1_41_MM7 | 3 | 5.81 | 0.42 | 0.610 | 0.087 |
| plsC_1_75_C | 0 | N.D. | N.D. | N.D. | N.D. |
| plsC_1_75_MM7 | 3 | 5.39 | 0.14 | 0.606 | 0.036 |
| pflB_1_47_C | 0 | N.D. | N.D. | N.D. | N.D. |
| pflB_1_47_MM7 | 3 | 5.63 | 0.24 | 0.662 | 0.015 |
| recA_1_77_C | 0 | N.D. | N.D. | N.D. | N.D. |
| recA_1_77_MM7 | 0 | N.D. | N.D. | N.D. | N.D. |
| ispF_1_32_C | 0 | N.D. | N.D. | N.D. | N.D. |
| ispF_1_32_MM7 | 0 | N.D. | N.D. | N.D. | N.D. |
| sucD_1_26_C | 0 | N.D. | N.D. | N.D. | N.D. |
| sucD_1_26_MM7 | 3 | 6.24 | 0.61 | 0.761 | 0.073 |
| yhbU_1_15_C | 0 | N.D. | N.D. | N.D. | N.D. |
| yhbU_1_15_MM7 | 3 | 5.71 | 0.37 | 0.570 | 0.017 |
| rpiA_1_58_C | 0 | N.D. | N.D. | N.D. | N.D. |
| rpiA_1_58_MM7 | 0 | N.D. | N.D. | N.D. | N.D. |
| tktB_1_15_C | 3 | 5.72 | 0.41 | 0.600 | 0.055 |
| tktB_1_15_MM7 | 3 | 5.64 | 0.36 | 0.570 | 0.049 |
| spoT_1_38_C | 3 | 4.82 | 0.41 | 0.583 | 0.019 |
| spoT_1_38_MM7 | 0 | N.D. | N.D. | N.D. | N.D. |
| ubiA_1_64_C | 3 | 5.10 | 0.74 | 0.536 | 0.223 |
| ubiA_1_64_MM7 | 3 | 5.46 | 0.29 | 0.675 | 0.055 |
| fadI_1_16_C | 0 | N.D. | N.D. | N.D. | N.D. |
| fadI_1_16_MM7 | 0 | N.D. | N.D. | N.D. | N.D. |
| clpP_1_34_C | 3 | 5.07 | 0.20 | 0.823 | 0.059 |
| clpP_1_34_MM7 | 3 | 5.48 | 0.48 | 0.623 | 0.073 |
| livJ_1_104_C | 3 | 5.26 | 0.04 | 0.585 | 0.034 |
| livJ_1_104_MM7 | 3 | 5.45 | 0.21 | 0.622 | 0.058 |
| pgpB_1_14_C | 3 | 6.35 | 0.14 | 0.607 | 0.086 |
| pgpB_1_14_MM7 | 0 | N.D. | N.D. | N.D. | N.D. |
| prpC_1_76_C | 3 | 6.62 | 0.27 | 0.602 | 0.009 |
| prpC_1_76_MM7 | 0 | N.D. | N.D. | N.D. | N.D. |
| sucB_1_24_C | 3 | 5.65 | 0.71 | 0.453 | 0.098 |
| sucB_1_24_MM7 | 3 | 5.06 | 0.70 | 0.334 | 0.084 |
| leuB_1_15_C | 3 | 5.93 | 0.65 | 0.611 | 0.051 |
| leuB_1_15_MM7 | 3 | 6.31 | 0.29 | 0.584 | 0.075 |
| idi_1_68_C | 3 | 6.11 | 0.66 | 0.341 | 0.044 |
| idi_1_68_MM7 | 0 | N.D. | N.D. | N.D. | N.D. |
| ispH_1_9_C | 0 | N.D. | N.D. | N.D. | N.D. |

| | | | | | |
|---|---|---|---|---|---|
| ispH_1_9_MM7 | 0 | N.D. | N.D. | N.D. | N.D. |
| elbB_1_121_C | 3 | 5.74 | 0.59 | 0.613 | 0.072 |
| elbB_1_121_MM7 | 3 | 5.30 | 0.41 | 0.621 | 0.027 |
| lon_1_7_C | 0 | N.D. | N.D. | N.D. | N.D. |
| lon_1_7_MM7 | 0 | N.D. | N.D. | N.D. | N.D. |
| dinG_1_11_C | 0 | N.D. | N.D. | N.D. | N.D. |
| dinG_1_11_MM7 | 3 | 6.13 | 0.42 | 0.615 | 0.055 |
| argB_1_109_C | 3 | 5.45 | 0.22 | 0.552 | 0.049 |
| argB_1_109_MM7 | 3 | 5.38 | 0.66 | 0.625 | 0.091 |
| pgpC_1_99_C | 0 | N.D. | N.D. | N.D. | N.D. |
| pgpC_1_99_MM7 | 0 | N.D. | N.D. | N.D. | N.D. |
| fbaA_1_28_C | 0 | N.D. | N.D. | N.D. | N.D. |
| fbaA_1_28_MM7 | 3 | 5.76 | 0.14 | 0.603 | 0.033 |
| talB_1_17_C | 3 | 3.75 | 0.66 | 0.575 | 0.125 |
| talB_1_17_MM7 | 3 | 5.39 | 0.25 | 0.616 | 0.030 |
| aroL_1_62_C | 0 | N.D. | N.D. | N.D. | N.D. |
| aroL_1_62_MM7 | 3 | 6.01 | 0.56 | 0.508 | 0.081 |
| ilvC_1_33_C | 3 | 5.82 | 0.33 | 0.625 | 0.034 |
| ilvC_1_33_MM7 | 3 | 5.73 | 0.43 | 0.647 | 0.102 |
| tktA_1_5_C | 0 | N.D. | N.D. | N.D. | N.D. |
| tktA_1_5_MM7 | 0 | N.D. | N.D. | N.D. | N.D. |
| leuA_1_29_C | 0 | N.D. | N.D. | N.D. | N.D. |
| leuA_1_29_MM7 | 3 | 5.85 | 0.23 | 0.547 | 0.015 |
| uvrB_1_37_C | 3 | 5.54 | 0.13 | 0.607 | 0.008 |
| uvrB_1_37_MM7 | 3 | 5.82 | 0.23 | 0.674 | 0.008 |
| ruvC_1_25_C | 3 | 6.05 | 0.51 | 0.480 | 0.089 |
| ruvC_1_25_MM7 | 3 | 6.61 | 0.56 | 0.462 | 0.090 |
| fabR_1_32_C | 3 | 5.81 | 0.38 | 0.546 | 0.022 |
| fabR_1_32_MM7 | 3 | 5.44 | 0.43 | 0.605 | 0.024 |
| yqeF_1_37_C | 0 | N.D. | N.D. | N.D. | N.D. |
| yqeF_1_37_MM7 | 0 | N.D. | N.D. | N.D. | N.D. |
| ispE_1_53_C | 3 | 5.59 | 0.31 | 0.218 | 0.113 |
| ispE_1_53_MM7 | 0 | N.D. | N.D. | N.D. | N.D. |
| clsB_1_64_C | 0 | N.D. | N.D. | N.D. | N.D. |
| clsB_1_64_MM7 | 3 | 5.73 | 0.13 | 0.630 | 0.029 |
| pta_1_24_C | 3 | 6.74 | 0.42 | 0.675 | 0.067 |
| pta_1_24_MM7 | 3 | 7.12 | 0.21 | 0.641 | 0.038 |
| fabB_1_24_C | 0 | N.D. | N.D. | N.D. | N.D. |
| fabB_1_24_MM7 | 3 | 5.20 | 1.25 | 0.579 | 0.083 |
| Non-targeting control | 3 | 6.24 | 0.60 | 0.580 | 0.083 |

| | | | | | |
|---|---|---|---|---|---|
| CRISPR control – dxs_1_17_C | 3 | 5.65 | 0.89 | 0.064 | 0.010 |
| WT MG1655::dCas9 | 4 | 5.10 | 0.48 | 0.005 | 0.002 |
| WT + pIpi | 4 | 5.12 | 0.19 | 0.675 | 0.077 |
| DD1 | 4 | 5.32 | 0.08 | 0.886 | 0.054 |
| DD2 | 4 | 5.43 | 0.09 | 0.780 | 0.008 |
| HW2e | 4 | 4.37 | 0.02 | 0.662 | 0.060 |
| HW2f | 4 | 4.46 | 0.17 | 0.671 | 0.041 |

**Table S4**. Average and standard deviation of OD600 and Lycopene yield (mg/(L*OD600)) for each identified sgRNA and reference strain (Related to Figure 4).

| Strain | Rep1 Yield (mg/L) | Rep2 Yield (mg/L) | Rep3 Yield (mg/L) | Avg Yield (mg/L) | SD Yield (mg/L) | p-value |
|---|---|---|---|---|---|---|
| accB_C | 1.27 | 2.07 | 3.16 | 2.17 | 0.95 | 0.213 |
| accB_MM7 | 1.73 | 2.09 | 2.28 | 2.03 | 0.28 | 0.008 |
| accA_C | 1.57 | 1.71 | 1.76 | 1.68 | 0.10 | 0.007 |
| accA_MM7 | 3.5 | 3.95 | 3.92 | 3.79 | 0.25 | 0.040 |
| accD_C | 1.64 | 2.67 | 2.56 | 2.29 | 0.57 | 0.105 |
| accD_MM7 | 1.82 | 2.49 | 2.9 | 2.40 | 0.55 | 0.130 |
| negC_rand_42 | 3.2 | 3.38 | 2.83 | 3.14 | 0.28 | 1.000 |

**Table S5**: Selected fatty acid biosynthesis genes from large scale sgRNA library. Average and standard deviations of lycopene yield (mg/L), as well as p-values from two-tailed, unequal variance, t-test for each strain versus NT control.
